# GFI1 as a novel regulator of γδ T cell development and the IL-17/IFNγ lineage commitment

**DOI:** 10.64898/2026.02.26.708021

**Authors:** Jennifer Fraszczak, David Obwegs, Kaifee Arman, Tanja Muralt, Bavanitha Thurairajah, Ève Mallet Gauthier, Irah L. King, Heather J. Melichar, Sagar, Tarik Möröy

## Abstract

GFI1 is a DNA-binding zinc finger transcription factor regulating the commitment of hematopoietic precursors to myeloid and lymphoid lineages. Here we report that GFI1 is expressed in γδ T cells and restricts the cellularity of RORγt^+^Vγ6^+^ γδT cells that produce high levels of IL-17A, while promoting the expansion of Vγ1^+^ and Vγ4^+^ γδT cells. Absence of GFI1 results in a pronounced bias toward the production of γδT17 cells, commencing post-birth. Additionally, we observe an expansion of RORγt^+^/MAF^+^ cells within the thymic DN1e population of GFI1-deficient mice. The DN1e population, along with other DN subsets in GFI1 knock-out (KO) mice, exhibits a distinctive γδT17 cell-specific transcriptomic profile. Specifically, DN1, DN3, and γδ T cells lacking GFI1 show upregulation of the B-ZIP transcription factor MAF, which regulates genes critical for γδT cells, such as *Il17a, Il22*, and *Blk* that are all induced in *Gfi1* deficient cells. The *Maf* gene is occupied by GFI1 in DN pre-T cells at cognate binding sites in its promoter region, suggesting that GFI1 acts as a direct repressor of *Maf*. We conclude that GFI1 functions as a novel regulator of Vγ6^+^ γδT17 precursor cells restricting their peripheral expansion by acting upstream of a MAF dependent regulatory network.

**Highlights:**

- GFI1 controls the expansion of γδ T cells in peripheral lymphoid organs and in barrier tissues.
- GFI1 specifically restricts expansion of γδ T cells secreting IL-17 by acting upstream of a MAF dependent transcriptional regulatory network.
- GFI1 plays a cell intrinsic role in controlling the generation of γδT cells through the repression of *Maf*.
- GFI1 controls a MAF^+^/RORγt^+^ γδT cell precursor cell population within the DN1e cell subset.

## Introduction

GFI1 is a 50kD transcription factor that contains an N-terminal SNAG domain and six C-terminal C_2_H_2_-type zinc fingers, which facilitate DNA binding to target genes^1–4^. The SNAG domain of GFI1 interacts with the histone demethylase KDM1A (LSD1) and recruits it with CoRest to target genes^5–8^. GFI1 and LSD1 are also part of the Nucleosome remodeling and deacetylase complex (NuRD)^9,10^, which enables transcriptional repression^5^. *Gfi1* was first identified in retrovirally induced murine T cell lymphoma^11,12^, but later, through the analysis of *Gfi1* deficient mice^13^, it was recognized that GFI1 controls the proliferation of hematopoietic stem cells^14^ as well as lymphoid/myeloid^15^ and monocyte/granulocyte lineage decisions in progenitor cells^2,16^. Moreover, GFI1 controls pre-T cell numbers in the thymus and the T cell differentiation process which begins at the early thymic progenitor (ETPs) stage. GFI1 is required for the survival and proliferation of the very early DN1 and DN2 pre-T cells and absence of GFI1 leads to the accumulation of an abnormal population between the DN1-DN2 stages, described as a block of differentiation^17^. In addition, GFI1 also plays a role in pre-TCR selection, where it regulates the proliferation and differentiation of DN3 cells into DN4 cells and is involved in major histocompatibility complex (MHC)-restricted positive and negative selection processes, highlighting its influence on the development of CD4 and CD8 single-positive T cells^17^. GFI1 has been shown to regulate pre-T cell differentiation through NOTCH1 signaling^18^, which is essential for the maintenance of ETPs and T cell lineage commitment and gene expression of lung Krüppel-like factor (KLF2) or inhibitor of DNA binding 1 and 2 (Id1 and Id2)^17^. GFI1 also regulates Foxo1 expression in thymic DP cells to maintain a specific transcriptional program in DP cells to assure the formation of mature T cells^19^. GFI1 also restrains the generation of regulatory T cells and plays a role in their functions^20–22^. In mature CD4^+^ T cells, GFI1 is induced to maintain an active Th2 program^22,23^ by either preventing the degradation of GATA3 or by blocking the production of IL-17 regulated by RORγt^24–28^. GFI1 also inhibits the Th1 program in activated CD4^+^ T cells^29^ and reins the Th9 polarisation^30^. In mature CD8^+^ T cells, GFI1 has recently been shown to control the formation of effector CD8^+^ T cells^31^ and the persistence of memory CD8^+^ T cells^32^ during infection.

To date, all studies on the function of GFI1 in T cell differentiation or peripheral T cell activity focused on the αβ T cell receptor-expressing populations, while a potential role for GFI1 in γδ T cells has not been investigated. Murine γδ T cells develop in waves starting in the fetal thymus at E14 with Vγ5^+^Vδ1^+^ dendritic epidermal T cells (DETC) followed by Vγ6^+^ and Vγ4^+^ γδ T cell subsets around fetal stage E16 homing to specific tissues like the lungs or the tongue^33^. Around birth, while DETC and Vγ6^+^ γδ T cell production declines drastically, new subsets of γδ T cells emerge that seed the intestine, liver, and peripheral lymphoid organs, respectively^34^. Different γδ T cell subsets have been identified^34–36^ but the two main subpopulations are γδT1 cells^37^, and γδT17 cells^38^. It is known that most γδT17 cells carry either the Vγ4 or the Vγ6 chain^54^, with Vγ6^+^ γδ T cells being produced almost exclusively at the fetal stage and the Vγ4^+^ γδ T cells at the fetal and perinatal stages^56–58^. Recent reports suggest that γδT17 and γδT1 cells can develop in the thymus *via* several pathways. One of these pathways is independent of the TCR and is initiated in two subsets of the DN1 thymic precursors called DN1d and DN1e^39,40,41^. From these subsets both IFNγ^+^ and IL17^+^ γδ T cells can emerge^41^. The second pathway to generate γδT17 and γδT1 cells in the thymus involves DN3 cells, where a strong TCR signal favors the IFNγ-producing lineage whereas a weak signal is associated with the development of γδT17 lineage^33,42–45^. Each γδ T cell subset has specific functions that are not only critical for the immune innate response against many pathogens^35,46^, but also play a role in autoimmune disorders or inflammation^47–51^. Moreover, clinical trials are also currently under way to test whether γδ T cells are effective in cancer immunotherapy^52^ creating a need to better understand their development and function. The role that transcription factors plays in establishing γδ T cells fate has been challenging and current knowledge remains limited.

Here we demonstrate that GFI1 is expressed in γδ T cells and that genetic ablation of GFI1 leads to a significant expansion of RORγt^+^Vγ6^+^γδT cells producing IL17 with a concomitant decrease in Vγ1^+^ and Vγ4^+^ γδT cells in adult mice. This bias toward the γδT17 subtype starts after birth in GFI1 knock-out (KO) mice and remains in place in the adult thymus, where a Vγ6^+^ subset secreting IL-17 becomes the predominant population among γδ T cells. In addition, GFI1 KO mice have an increased number of c-MAF^+^/RORγt^+^ cells in the DN1e subset, which has been described to contain precursors for γδT17 cells^40,41^. *Gfi1*-deficient DN thymocytes and γδ T cells show an upregulation of the B-Zip transcription factor c-MAF, which is known to regulate the production of γδT17 cells suggesting that GFI1 controls the cellularity of γδ T cells and their progenitors though the regulation of c-MAF.

## Results

### *Gfi1* deficient mice show an expansion of γδ T cells in barrier tissues and in peripheral lymphoid organs

To gain more insight into a potential role of GFI1 in the immune compartment of barrier tissues, we performed single cell RNA-sequencing (scRNA-seq) of lung cells from GFI1 WT and KO mice. We noticed that the UMAP representation showed two clusters (clusters 5 and 8) of *Cd4^-^*, *Cd8a*^-^, *Cd8b*^-^, *Trdc*^+^ cells within the *Cd3e*^+^ lymphocyte compartment that were overrepresented in GFI1 KO mice (**Fig. 1A-C & Suppl. Fig.1A & 1B**). These two clusters were annotated as γδ T cells while all other clusters were conventional CD4^+^ or CD8^+^ T cells (**Fig. 1A-C & Suppl. Fig.1A & 1B**). Flow cytometry confirmed the presence of an increased proportion of γδ T cells in the lungs, the spleen, lymph nodes (LN) as well as in the intestine lamina propria of GFI1 KO mice (**Fig. 1D-F** & **Suppl. Fig. 1C**). This observation stood in stark contrast to the numbers of CD4^+^ T cells, which are known to be reduced in *Gfi1* deficient mice^17,18,53^. Accordingly, the proportion of cells from GFI1 KO mice belonging to the CD4^+^ T cell clusters (clusters 7, 10 and 0) were reduced compared to WT mice (**Fig. 1A-C & Suppl. Fig.1A & 1B**), which was also seen by flow cytometry (**Suppl. Fig. 1D**). The same observations were made with *vav-cre GFI1^flox/flox^* mice where GFI1 is deleted solely in hematopoietic cells. As in the germline GFI1 KO mice, γδ T cells from *vav-cre GFI1^flox/flox^* mice were expanded in number and frequency compared to controls (**Fig. 1 G-I**), while CD4^+^ T cells were reduced (**Suppl. Fig. 1E)**. This excluded the non-hematopoietic environment as a potential cause for γδ T cell expansion in GFI1 KO mice.

**Fig. 1:**
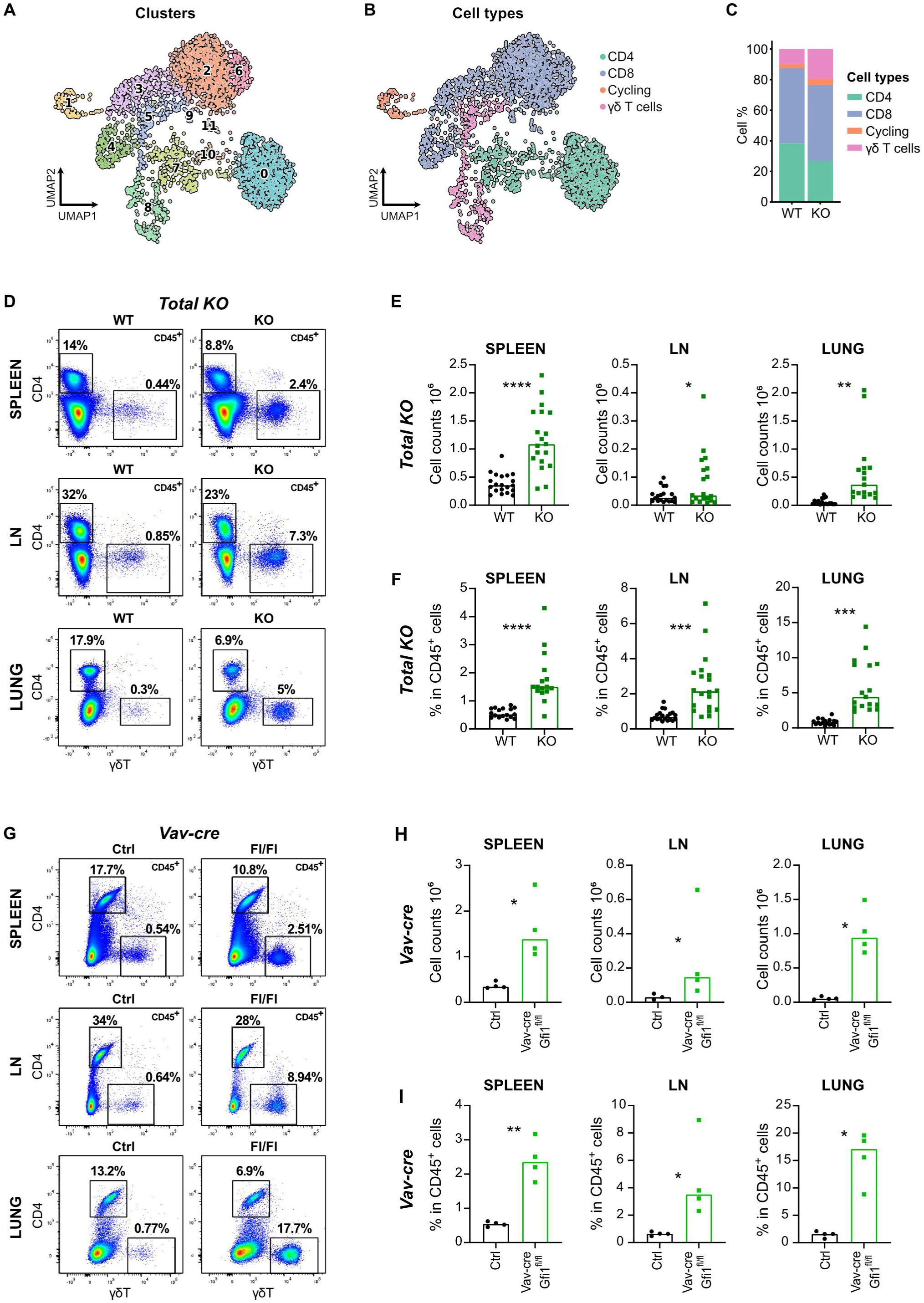
Phenotype of γδ T cells in adult GFI1 KO mice. **A** & **B**: UMAP representation based on gene expression profiling by sc-RNA-seq depicting 11 clusters in the CD3^+^ lymphocyte compartment in the lung tissue of WT and GFI1 KO mice (**A**) and the defined subsets labeled in different colors (**B**). **C**: Bar plots showing the distribution of the defined populations in the indicated genotypes. **D**: Representative plots of lung, splenic and LN cells from WT and GFI1 KO mice analyzed by flow cytometry and stained to identify γδ T cells (CD45^+^CD4^-^γδT^+^) after gating on live cells (according to the FSC/SSC) and singlets. **E & F**: Cell numbers (**E**) and frequencies in CD45^+^ cells (**F**) of γδ T cells (CD4^-^γδT^+^) in the lung, spleen and LN of WT and GFI1 KO mice. **G**: Representative plots of lung, splenic and LN cells from control and *vav-cre GFI1^flox/flox^* mice analyzed by flow cytometry and stained to identify γδ T cells (CD45^+^CD4^-^γδT^+^) after gating on live cells (according to the FSC/SSC) and singlets. **H & I**: Cell numbers (**H**) and frequencies in CD45^+^ cells (**I**) of γδ T cells (CD4^-^γδT^+^) in the lung, spleen and LN of control and *vav-cre GFI1^flox/flox^* mice. Adult mice meaning mice older than 5 weeks. Statistical testing was performed with paired *t*-test using Prism v9 (GraphPad). *p*-Values are shown as * < 0.05, ** < 0.01, *** < 0.005, and **** < 0.001 where each statistical significance was found.

### Specific expansion of **γδ**T17 cells in GFI KO mice

To determine if the expansion of γδ T cells in GFI1 KO mice was specific to a γδ T cells subtype, we further examined the key features of expanded γδ T cells in GFI1 KO lungs. Expanded lung γδ T cells showed transcriptomic features of γδT17 cells indicated by the expression of *Blk*, *Maf*, *Rorc* and *Scart*, in particular cluster 8 (**Fig. 2A**). We analysed these cells further and found that a large proportion of the expanded γδ T compartment in GFI1 KO lacked CD27 (**Fig. 2B, Suppl. Fig. 2A**), a marker associated with the γδT1 subtype^54^. Consistent with this observation, we also found that up to 50% of *Gfi1* deficient γδ T cells are RORγt^+^ in the lung, spleen, and LN (**Fig. 2C-E**). Similar data were obtained with *vav-cre GFI1^flox/flox^*mice where increased frequencies and numbers of RORγt^+^ γδ T cells were also evident compared to controls (**Suppl. Fig. 2B-D**). The upregulation of RORγt was specifically seen in γδ T cells and was not found in *Gfi1*-deficient CD4^+^ T cells (**Fig. 2C, Suppl. Fig. 2B**).

**Fig. 2:**
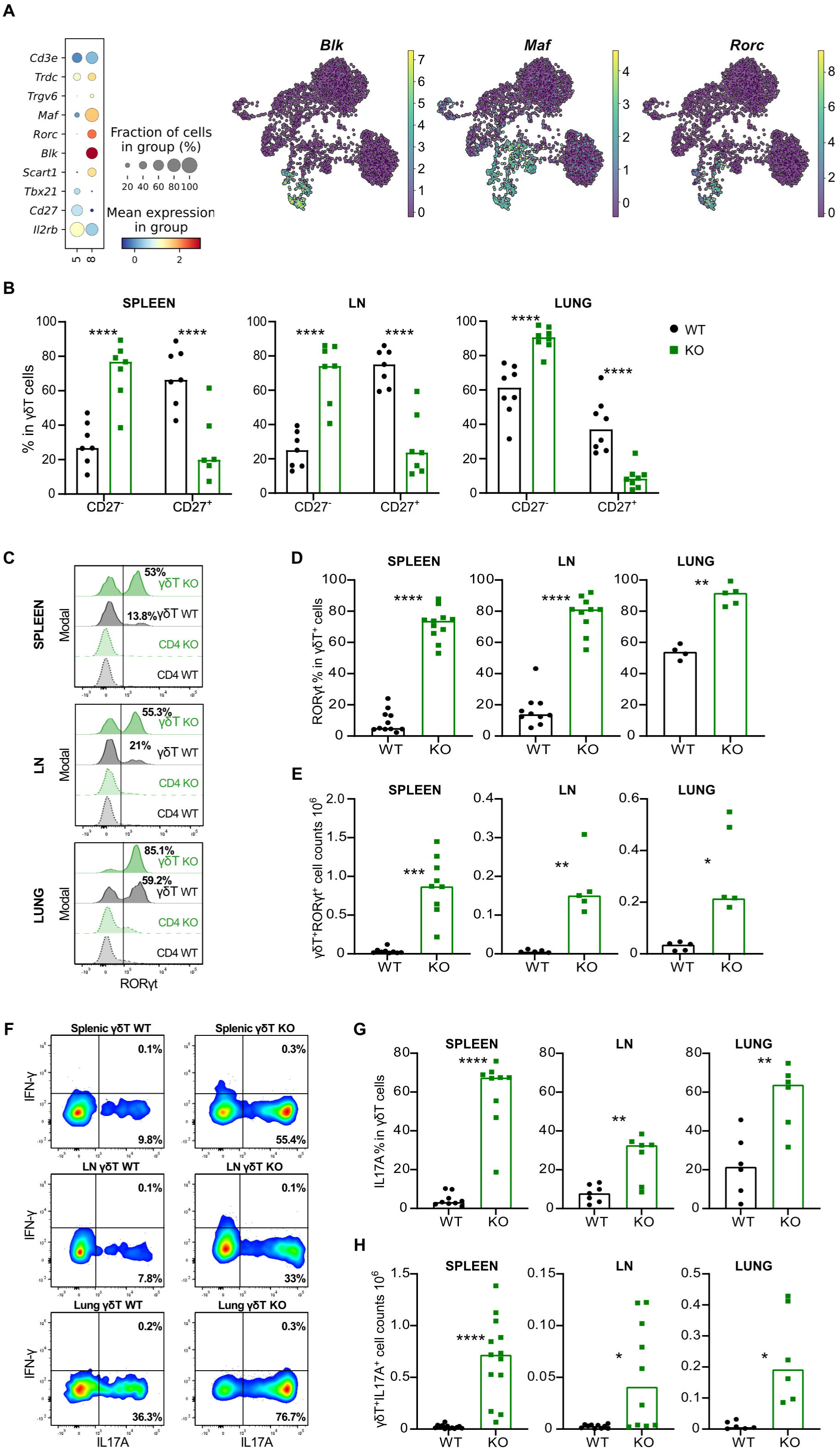
**Expansion of the ROR**γ**t^+^Il-17a^+^** γδ**T subset in adult GFI1 KO mice. A:** Dot plot showing the key differentially expressed genes in cluster 5 and 8 and UMAP representation showing expression of *Blk, Maf and Rorc* genes in the CD3^+^ lymphocyte compartment in the lung tissue of WT and GFI1 KO mice. Color represents the normalized expression of the gene. **B:** Frequencies of CD27^+^ and CD27^-^ cells in splenic, LN and lung CD45^+^CD4^-^γδT^+^ from WT and GFI1 KO mice. **C**: Representative normalized histogram of RORγt^+^ cells in either splenic, LN or lung CD45^+^CD4^-^γδT^+^ and CD45^+^CD4^+^γδT^-^ from WT and GFI1 KO mice. **D**: Frequencies of RORγt^+^ cells in splenic, LN and lung CD45^+^CD4^-^γδT^+^ comparing WT and GFI1KO mice. **E**: Splenic, LN and lung CD45^+^CD4^-^γδT^+^RORγt^+^ cell counts in WT and GFI1 KO mice. **F**: Representative plots of lung, splenic and LN cells from WT and KO mice treated with PMA, ionomycin and Golgi Plug for 3.5 hours and analysed by flow cytometry. The cells were gated on singlets CD45^+^CD4^-^γδT^+^ before gating for IFNγ and IL17A. **G**: Frequencies of IL17A^+^ in splenic, LN and lung CD45^+^CD4^-^ γδT^+^ comparing WT and GFI1 KO mice. **H**: Cell numbers of splenic, LN and lung CD45^+^CD4^-^γδT^+^IL17A^+^ in WT and GFI1 KO mice. Adult mice meaning mice older than 5 weeks. Statistical testing was performed with multiple paired *t*-test for **B** and paired *t*-test for **D-E** and **G-H** using Prism v9 (GraphPad). *p*-Values are shown as * < 0.05, ** < 0.01, *** < 0.005 and **** < 0.001 where each statistical significance was found.

To test the functionalities of GFI1 KO γδ T cells, we cultured cells from the spleen, LN, and lung of WT and GFI1 KO mice and treated them with PMA and ionomycin in the presence of Golgi plug. We found that large proportions of *Gfi1* deficient γδ T cells from spleen, LN and lungs produced IL17A (**Fig. 2F & 2G**), which led to significantly increased numbers of IL17A^+^ γδ T cells in the KO culture when compared to WT controls (**Fig. 2H**). Moreover, GFI1 KO γδ T cells were also able to produce IL22 (**Suppl. Fig. 2E-G**), a cytokine often produced together with IL17A by γδT17 cells^55^. These findings suggest that the absence of GFI1 leads specifically to the expansion of γδT17 cells, which produce IL17 after stimulation.

### V**γ**6-expressing **γδ**T17 cells expand in the absence of GFI1

To characterize further the subtype of γδT17 cells, we performed a scRNA-seq analysis on sorted γδ T cells from spleens of WT and GFI1 KO mice. We annotated the clusters based on our previously published study^59^, and 14 clusters were identified in splenic γδ T cells (**Fig. 3A & Suppl. Fig. 3A**), with 6 clusters (0, 1, 3, 4, 5 and 12) expressing genes associated with the Vγ6^+^ γδ T cell subtype such as *Trgv6*, *Cxcr6, Scart1*, and *Rorc*^60^ (**Fig. 3B & 3C**). These 6 clusters represented almost the totality of the splenic γδ T cells sorted from GFI1 KO mice (**Fig. 3C-E**). Consistent with the previous observations, we found that the most upregulated genes in GFI1 KO splenic γδ T cells compared to WT controls were associated with the mature Vγ6^+^ γδT17 cells like *Scart1*, *Maf*, *Blk* and *Ccr2*^60^, while genes associated with exchanging and adaptive-like γδ T cells such as *Tcf7* and *Lef1* were downregulated (**Fig. 3F**), which was in agreement with the expansion of Vγ6^+^ γδ T cells in the spleen and in the lung of GFI1 KO mice compared to controls determined by flow cytometry (**Fig. 3G & 3H**).

**Fig. 3:**
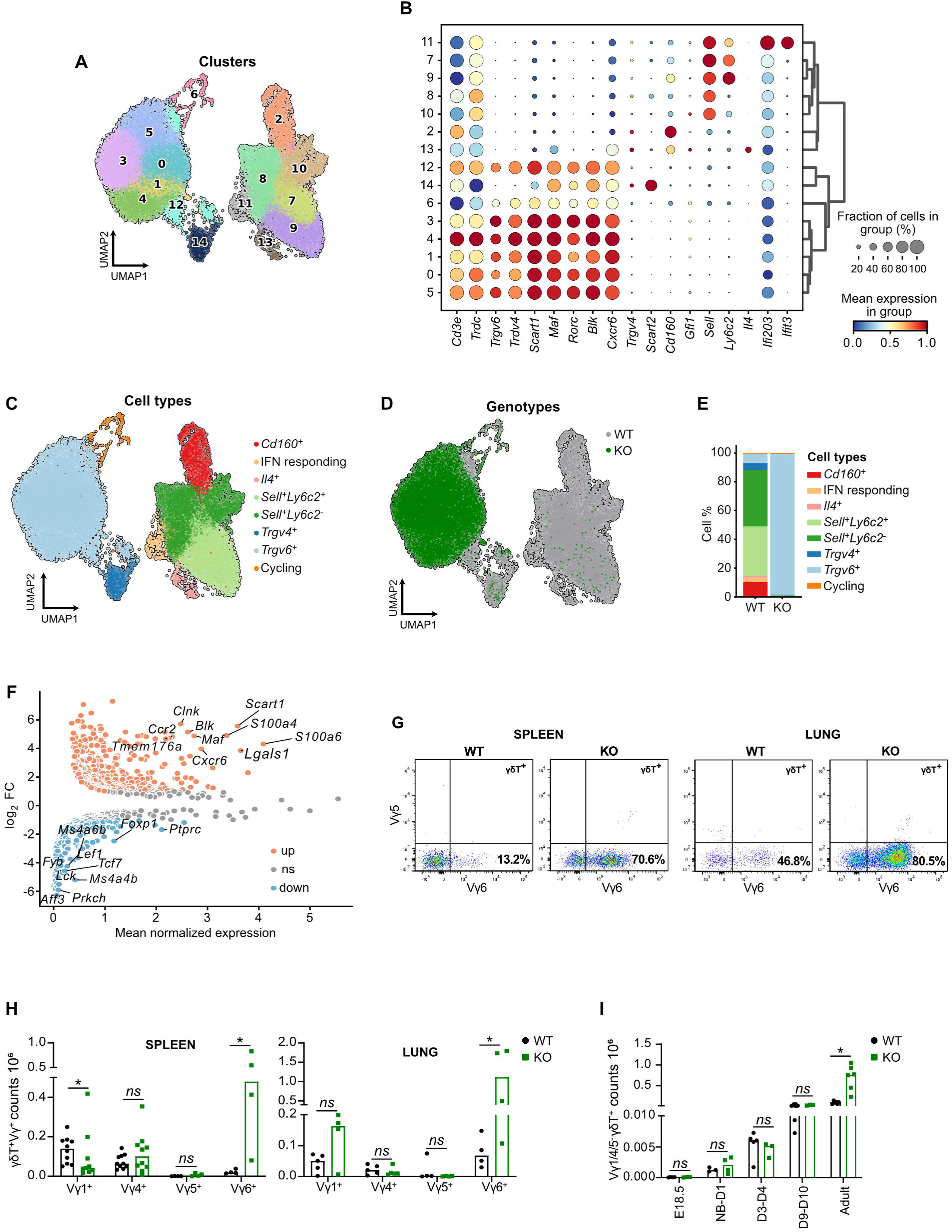
Expansion of. **V**γ**6^+^** γδ**T17 cells in the periphery of adult GFI1 KO mice. A**: UMAP representation based on gene expression profiling by sc-RNA-seq depicting 14 clusters in sorted splenic γδ T cells of WT and GFI1 KO mice. **B**: Dot plot showing the key differentially expressed genes in each cluster shown in Fig. 3A. Color represents the scaled mean expression of the gene in the respective cluster, and dot size represents the fraction of cells in the cluster expressing the gene. **C** and **D**: UMAP representation of sorted splenic γδ T cells of WT and GFI1 KO mice colored by cell type (**C**) or genotype (**D**). **E**: Bar plots showing the distribution of cell types found in splenic γδ T cells in the indicated genotypes. **F**: Differential gene expression showing key up-or down-regulated genes in GFI1 KO splenic γδ T cells versus WT splenic γδ T cells. **G**: Representative plots of Vγ5^+^ and Vγ6^+^ in γδT^+^ cells from WT and GFI1 KO spleen and lung after gating on live cells (according to FSC/SSC) and singlets. **H**: Vγ1^+^, Vγ4^+^, Vγ5^+^ or Vγ6^+^ γδT^+^ cell counts in the spleen and lung of adult WT and GFI1 KO mice. **I**: Cell numbers of Vγ1^-^Vγ4^-^Vγ5^-^ γδT^+^ cells at different time point in WT and GFI1 KO spleen, with adult meaning mice older than 5 weeks. Adult mice meaning mice older than 5 weeks. Statistical testing was performed with multiple Mann-Whitney tests using Prism v9 (GraphPad). *p*-Values are shown as * < 0.05, ** < 0.01, *** < 0.005, and **** < 0.001 where each statistical significance was found.

Stimulation of sorted splenic Vγ1^-^Vγ4^-^Vγ5^-^ γδ T cells (i.e.mostly Vγ6) with either IL23 or IL7 also showed that cells from the GFI1 KO mice were more responsive than WT cells as shown by the higher production of IL17A or the increased phosphorylation of STAT3 (**Suppl. Fig. 3B**). This indicated that most of the cells from GFI1 KO mice were mostly constituted of functional Vγ6^+^ γδ T cells, capable of responding to cytokine stimulation. Unexpectedly, the expansion of Vγ6^+^ γδ T cells in the spleen started only after birth (**Fig. 3I**), at a point in development when the generation of this subtype is not supposed to be maintained, suggesting that GFI1 controls and possibly restricts, the expansion of functional Vγ6^+^ γδT17 cells only in adult mice in the periphery.

### Defects in **γδ** T cell production are detected in the thymus of adult mice

To determine whether the γδ T cell phenotype observed in GFI1 KO mice was due to a developmental defect, we analyzed γδ T cells from the thymus, where γδ T cells originate and diversify^61^. Even though, absolute numbers of γδ T cells were similar in GFI1 KO and WT adult thymus, both numbers and frequencies of IL17A^+^/RORγt^+^ cells were significantly increased in *Gfi1* deficient mice compared to WT controls (**Fig. 4A-F**). Again, similar data were obtained with *vav-cre Gfi1^fl/fl^* mice (**Suppl. Fig. 4A**-**C**). scRNA-seq analysis of sorted thymic γδ T cells from WT and GFI1 KO mice revealed 14 clusters with 5 clusters (clusters 6, 9, 10, 11 and 13) expressing genes specific to the Vγ6^+^ γδT17 subtype such as *Scart1, Rorc, Maf* and *Trgv6* (**Fig. 4G-4I and Suppl. Fig. 4D**)^60,62^. Clusters 6, 9, 10, 11 and 13 form a larger cluster containing only Vγ6^+^ γδT17 cells identified as *Trgv6^+^* and cluster 14 was annotated as *Gzma^-^Scart2^+^*and could represent a cluster of Vγ4^+^ γδT17 cells based on the expression of *Maf, Rorc and Scart2* (**Fig. 4H&I**). The low expression of Vγ4 is consistent with limitations in 3’ sequencing protocols used here, which don’t capture TCR variable regions reliably.

**Fig. 4:**
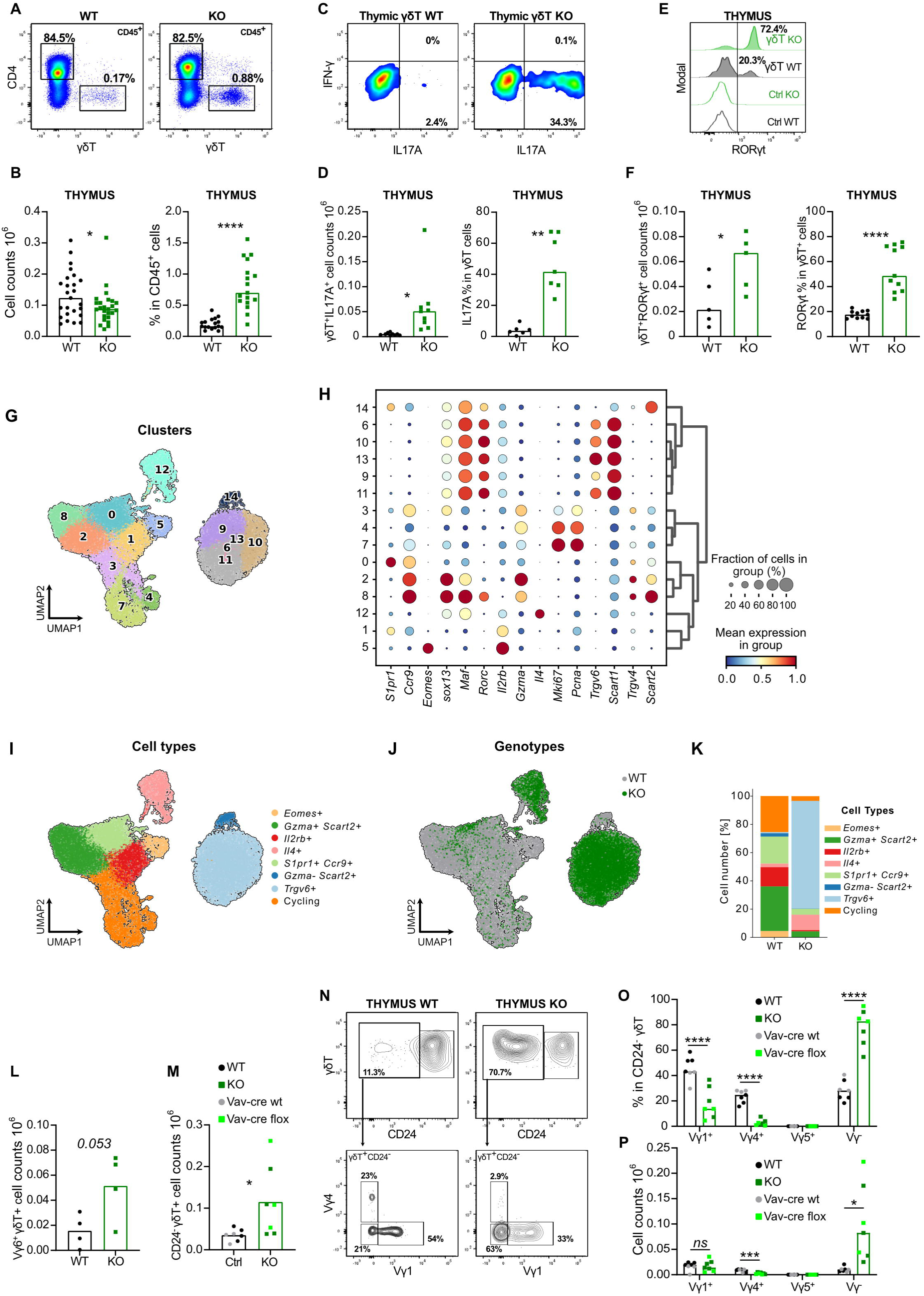
Expansion of. **V**γ**6^+^** γδ**T17 cells in the thymus of adult GFI1 KO mice. A:** Representative plots of thymic CD45^+^CD4^-^γδT^+^ cells in WT and GFI1 KO mice after gating on live cells (according to FSC/SSC) and singlets. **B**: γδ T cell counts (CD45^+^CD4^-^γδT^+^, left) and frequencies in CD45^+^ cells (CD4^-^γδT^+^, right) in WT and GFI1 KO thymus. **C**: Representative plots of thymic γδ T cells (CD45^+^CD4^-^γδT^+^) treated with PMA, ionomycin and Golgi Plug for 3.5 hours and stained for IL17A and IFNγ. The cells were gated on singlets CD45^+^CD4^-^γδT^+^ before gating for IFNγ and IL17A. **D**: CD45^+^CD4^-^γδT^+^IL17^+^ cell counts (left) and IL17A^+^ cell frequencies in CD45^+^CD4^-^γδT^+^ (right) in WT and GFI1 KO thymus. **E**: Representative normalised histogram for RORγt and control isotype (Ctrl) in CD45^+^CD4^-^γδT^+^ from WT and GFI1 KO thymic cells. **F**: CD45^+^CD4^-^γδT^+^RORγt^+^ cell counts (left) and RORγt^+^ cell frequencies in CD45^+^CD4^-^γδT^+^ (right) in WT and GFI1 KO thymus. **G**: UMAP representation based on gene expression profiling depicting 14 clusters in sorted thymic γδ T cells of WT and GFI1 KO mice. **H**: Dot plot showing the key differentially expressed genes in each cluster shown in Fig. 4G. Color represents the scaled mean expression of the gene in the respective cluster, and dot size represents the fraction of cells in the cluster expressing the gene. **I** and **J**: UMAP representation of sorted thymic γδ T cells of WT and GFI1 KO mice colored by cell type (**I**) or genotype (**J**). **K**: Bar plots showing the distribution of each cell type found in thymic γδ T cells in the indicated genotypes. **L**: Thymic Vγ6^+^ γδT^+^ cell counts in adult WT and GFI1 KO mice based on cytometry staining. **M**: Cell numbers of CD24^-^ γδT^+^ in adult thymus of WT and GFI1 KO mice or control (*vav-cre GF1^wt/wt^*) and *vav-cre GFI1^flox/flox^*mice. **N**: Representative plots of CD24 expression in thymic γδ T cells and Vγ chains expression in thymic CD24^-^ γδ T cells from adult WT and GFI1 KO mice. **O & P**: Vγ chain frequencies (**O**) and Vγ chain cell counts (**P**) in thymic CD24^-^ γδ T cells from adult WT and GFI1 KO mice or control (*vav-cre GF1^wt/wt^*) and *vav-cre GFI1^flox/flox^*mice. Adult mice meaning mice older than 5 weeks. Statistical testing was performed with paired *t*-test or multiple paired *t*-test using Prism v9 (GraphPad). *p*-Values are shown as * < 0.05, ** < 0.01, *** < 0.005, and **** < 0.001 where each statistical significance was found.

Interestingly, the cells found in the *Trgv6^+^* cluster were exclusively from GFI1 KO mice and represented the majority of its thymic γδ T population while the *Trgv4^+^*cluster was only found in WT thymus (**Fig. 4J&K**). Consistent with this data, the most upregulated genes in GFI1 KO thymic γδ T cells were again genes associated to the Vγ6 subset such as *Scart1* and *Cxcr6* (**Suppl. Fig. 4E**), which is consistent with the over representation of the of Vγ6 γδT17 subtype already seen in the thymus of GFI1 KO mice. This bias towards the Vγ6 γδ T subset in adult GFI1 KO thymus was also confirmed by flow cytometry by using the specific antibody against the Vγ6 chain showing a strong tendency to higher numbers of Vγ6^+^ γδ T cells found in adult thymocytes lacking GFI1 compared to WT thymus (**Fig. 4L**). However, when γδ T cells numbers that are CD24^-^ are enumerated in adult thymocytes lacking GFI1, the difference to control mice becomes significant (**Fig. 4M, N**). Including a CD24 marker also rendered the gating on Vγ chain populations more obvious in *Gfi1* deficient γδ T cells (**Fig. 4N**). Again, we could confirm a higher proportion and cell count of Vγ1^-^Vγ4^-^Vγ5^-^ (most likely meaning Vγ6^+^) cells in CD24^-^γδ T cells lacking GFI1 in adult mice (**Fig. 4N-P**). The bias was also confirmed by staining for CD44 and CD45RB^33,43^, known to distinguish the different subsets of γδ T cells, with significantly increased numbers of CD44^+^CD45RB^-^ cells (known as γδT17) within the CD24^-^ γδ T cell subset in GFI1 KO thymus (**Suppl. Fig. 4F, G**). As observed in the spleen and lungs, the over representation of Vγ6^+^ γδ T cells in the GFI1 KO thymus also started after birth (**Suppl. Fig. 4H-K**) with higher frequencies of Vγ1^-^Vγ4^-^Vγ5^-^ (most likely meaning Vγ6^+^) cells in CD24^-^γδ T cells of the GFI1 KO young mice and a slight but significant increased number of this population at day 8-10, also seen at D18-20 after birth. This data suggested that the absence of GFI1 leads to a γδ T cell developmental defect with biased production of Vγ6^+^ γδ T cells over the other γδ T cell subtypes.

### GFI1 plays a cell intrinsic role in **γδ**T cells

To determine whether the loss of GFI1 can directly impact γδ T cell development, we evaluated GFI1 expression levels in these cells. First, flow cytometric analyses of cells from a GFI1 reporter mouse which carries an EGFP cDNA “knock-in” in the *Gfi1* locus placing it under the control of the *Gfi1* promoter^63^ indicated that γδ T cells from spleen, LN, thymus, and lung express higher levels of *Gfi1* than conventional CD4^+^ T cells (**Fig. 5A**). Additionally, a higher expression of *Gfi1* mRNA in sorted splenic and lung γδ T cells compared to naïve CD4^+^ T cells, where GFI1 is poorly expressed^53,63^ was also confirmed by q-RT-PCR (**Fig. 5B**). *Gfi1* expression in thymic γδ T cells was even higher than in DN1 but lower than in DN3 pre-T cells (**Fig. 5C**), while *Gfi1b* mRNA, which was used as a negative control, was not detected in γδ T cells (**Suppl. Fig. 5A**).

**Fig. 5:**
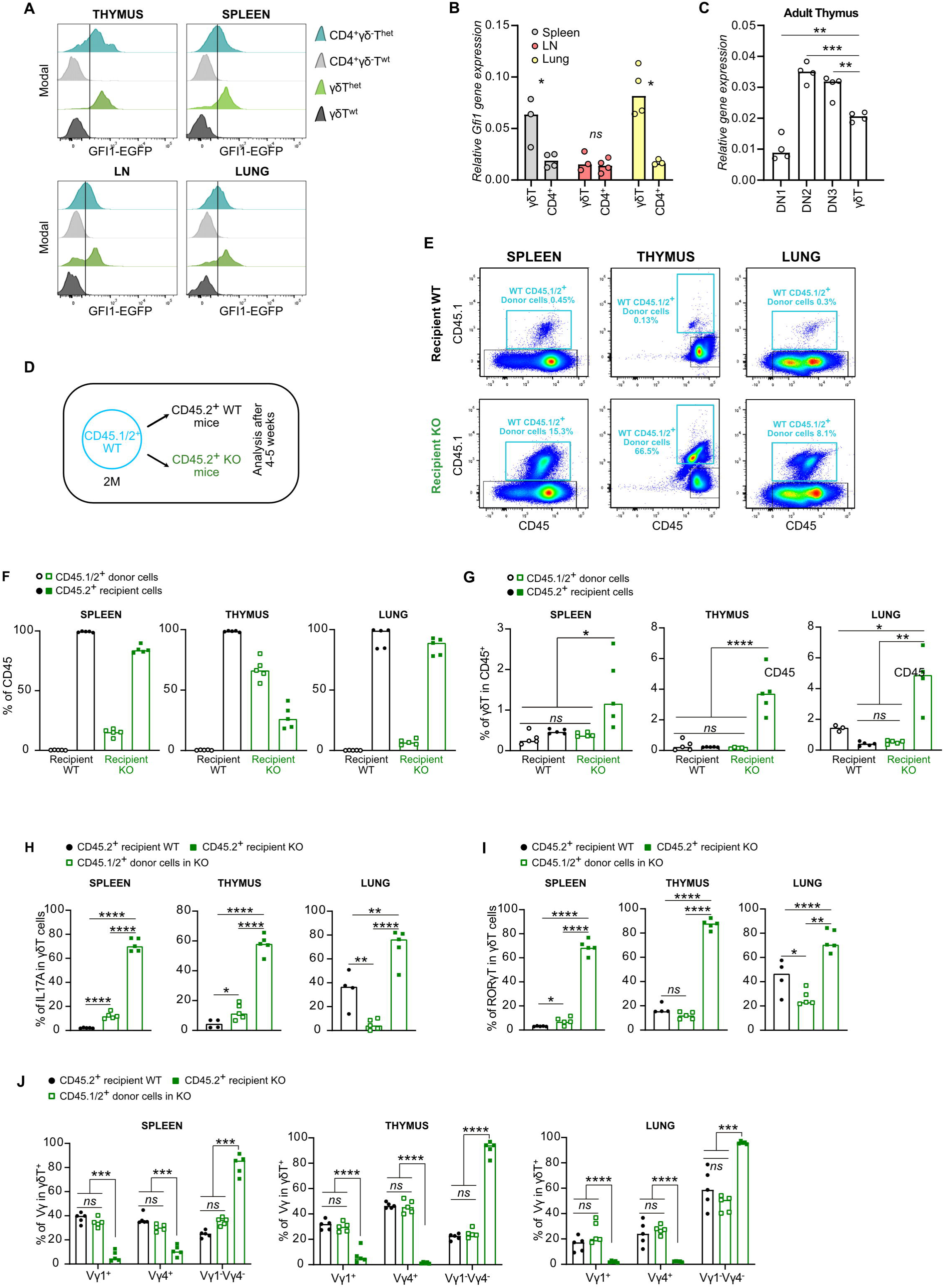
GFI1 intrinsically controls the generation of γδ. T cells. **A:** GFP^+^ cells indicating *Gfi1* expression in either γδ T cells (CD45^+^CD4^-^γδT^+^) or CD4^+^ T cells (CD45^+^CD4^+^γδT^-^) from spleen, LN, lung, and thymus from GFI1:GFP reporter mice^63^ (het) and controls (wt). **B** & **C**: Relative *Gfi1* expression in γδ T and CD4^+^ T cells from Spleen, LN and Lung (**B**) and in thymic γδ T, DN1, DN2 and DN3 cells (**C**) measured by RT-PCR. *Gapdh* was used as a housekeeping gene. **D**: Scheme representing the transplantation strategy of CD45.1/2^+^ WT BM cells into either CD45.2^+^ WT or GFI1 KO recipient mice. **E**: Representative plots showing the CD45.1/2^+^ donor cells in the spleen, thymus and lung of non-irradiated WT and KO recipients. **F**: Frequencies measured by flow cytometry of either WT CD45.1/2^+^ donor cells or CD45.2^+^ recipient cells in the spleen, lung and thymus of transplanted WT and GFI1 KO recipient mice. **G**: Frequencies measured by flow cytometry of either WT donor γδ T cells (in CD45.1/2^+^ cells) or endogenous recipient γδ T cells (in CD45.2^+^ cells) in the spleen, lung and thymus of transplanted WT and GFI1 KO recipient mice. **H**: IL17A frequencies in either donor WT γδ T cells (CD45.1/2^+^ cells) or endogenous recipient γδ T cells (in CD45.2^+^ WT or in CD45.2^+^ GFI1 KO cells) in the spleen, lung and thymus of transplanted WT and GFI1 KO recipient mice. **I**: RORγt frequencies in either WT donor γδ T cells (CD45.1/2^+^ cells) or endogenous recipient γδ T cells (in CD45.2^+^ WT or in CD45.2^+^ GFI1 KO cells) in the spleen, lung and thymus of transplanted WT and GFI1 KO recipient mice. **J**: Vγ1^-^Vγ4^-^ frequencies in either WT donor γδ T cells (CD45.1/2^+^ cells) or endogenous recipient γδ T cells (in CD45.2^+^ WT or in CD45.2^+^ GFI1 KO cells) in the spleen, thymus and lung of transplanted WT and GFI1 KO recipient mice. Statistical testing was performed with multiple paired *t*-test using Prism v9 (GraphPad). *p*-Values are shown as * < 0.05, ** < 0.01, *** < 0.005, and **** < 0.001 where each statistical significance was found.

Next, we wished to determine whether the bias towards the γδT17 subtype in GFI1 KO mice was due to the intrinsic lack of GFI1 in these cells or a consequence of the specific environment created by the absence of GFI1 in the hematopoietic compartment. To clarify this, we transplanted CD45.1/2^+^ WT bone marrow (BM) cells^64,65^ into non-irradiated CD45.2^+^ GFI1 KO mice and analysed the animals after 4-5 weeks. This enabled us to study the impact of the GFI1 KO environment on WT γδ T cells generated during adulthood (**Fig. 5D**). We were able to identify a small pool (between 15-20%) of WT hematopoietic cells identified as CD45.1/2^+^ cells amongst CD45.2^+^ GFI1 KO deficient cells representing 80-85% of the hematopoietic cells in the spleen and lung of the recipients (**Fig. 5E & 5F**). As a control, we also transplanted CD45.1/2^+^ WT BM cells into non-irradiated CD45.2^+^ WT recipient mice. However, as expected, the frequencies of CD45.1/2^+^ WT donor cells in non-irradiated CD45.2^+^ WT recipients were very low in the organs analysed (**Fig. 5E & 5F**). The frequencies of WT donor splenic γδ T cells in GFI1 KO and WT recipients were similar, while the frequencies of the endogenous GFI1 KO host γδ T cells were still elevated compared to WT donor or recipient γδ T cells (**Fig. 5G and Suppl. Fig. 5B**). We obtained the same results in lungs and thymus of the GFI1 KO recipient mice when they were transplanted with WT BM cells (**Fig. 5G and Suppl. Fig. 5B**). We found that the splenic, thymic, and lung WT γδ T cells in the KO recipient mice did not show an overexpression of RORγt or overproduction of IL17A when compared to the endogenous GFI1 KO host γδ T cells (**Fig. 5H, I and Suppl. Fig. 5C, D**), even though a slight increase of IL17A and RORγt could be seen compared to recipient WT γδ T cells in the spleen (**Fig. 5H, I and Suppl. Fig. 5C, D**). Moreover, the distribution of the different Vγ chain subtypes was similar between the WT γδ T donors in the GFI1 KO recipient and the WT γδ T cells in all analysed organs (**Fig. 5J and Suppl. Fig. 5E**). This suggested that the GFI1 KO environment has no or very little effect on γδ T cells and cannot explain the preferential bias toward the γδT producing IL17 in *Gfi1* deficient mice.

### Identification of a potentially precursor population in adult GFI1 KO mice

γδ T cells are thought to differentiate from progenitors located in the thymic DN compartment, which includes DN1, DN2, DN3 and DN4 subsets^33^. We set out to determine whether DN cells from GFI1 KO mice may have gained the ability to preferentially produce Vγ6^+^ γδ T cells. To test this, we cultured total DN cells from adult WT and GFI1 KO thymi on OP9-DL1 cells (**Suppl. Fig. 6A**), by excluding γδ T cells, NK cells, myeloid cells, B cells, and conventional T cells (**Suppl. Fig. 6B, C)**. Under these conditions, we observed that GFI1 KO DN cells can give rise to higher proportions of the CD24^-^CD44^+^CD45RB^-^ γδ T cells (known as γδT17) in the CD45^+^ cells compared to WT DN cells (**Fig. 6A, B**). These data suggested that the bias towards the γδT17 subset in the absence of GFI1 has its origin in early precursors of γδ T cell differentiation, which are not yet fully committed to the γδ T cell lineage.

**Fig. 6:**
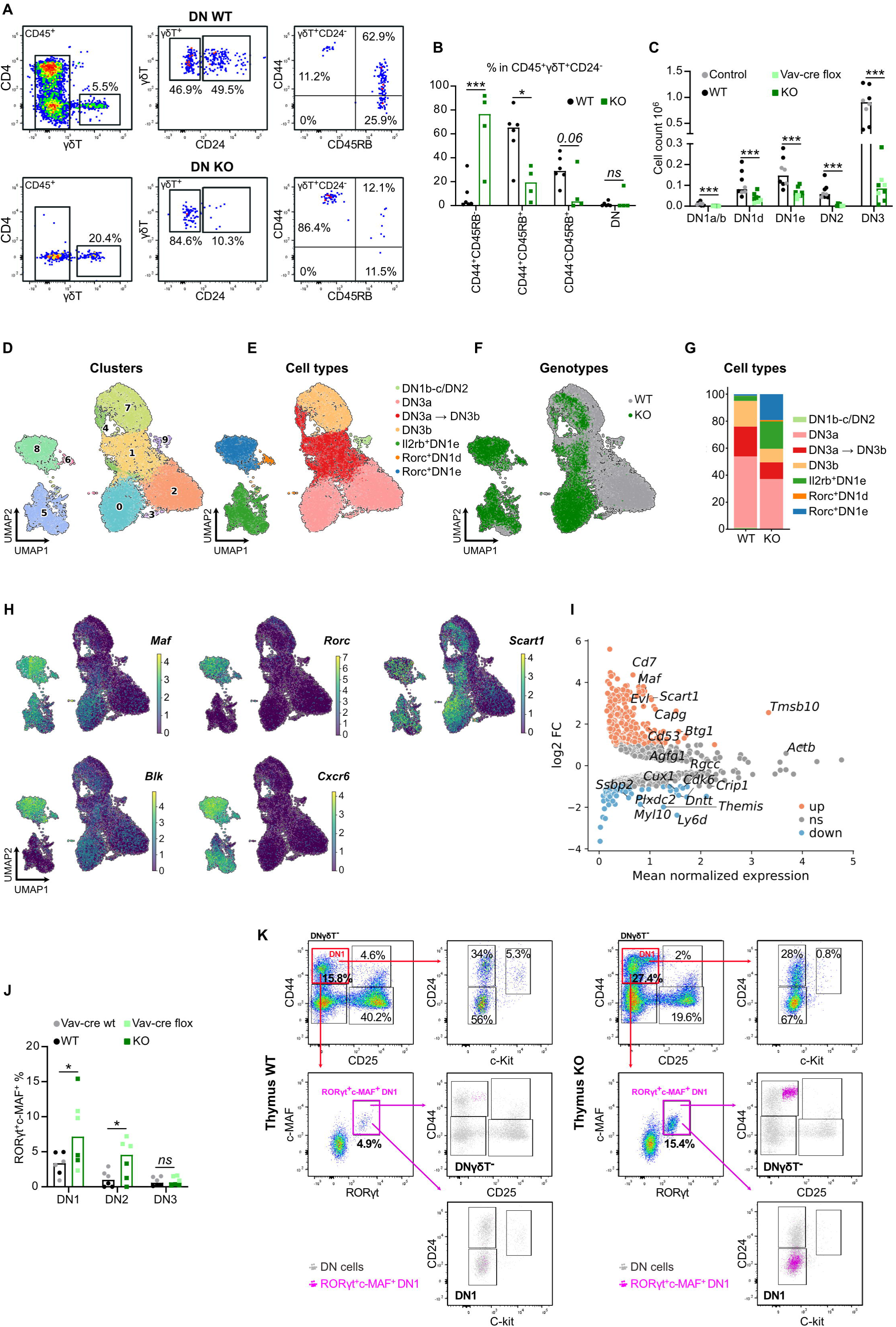
Precursors in thymus of adult mice A: Representative plots of WT or GFI1 KO thymic DN progenitors cultured on OP9-DL1 in the presence of FLT3 and IL7 for 12-14 days. After gating on singlets CD45^+^γδT^+^CD24^-^ cells, mature CD24^-^ γδT subsets were distinguished according to the expression of CD44 and CD45RB. **B**: Frequencies of each indicated subset from Fig. 6A. **C**: DN cell counts in the thymus of WT and GFI1 KO mice or control (*vav-cre GF1^wt/wt^*) and *vav-cre GFI1^flox/flox^*mice. DN1a/b cells were characterized as CD4^-^CD8^-^γδT^-^CD44^+^CD25^-^CD24^+^Ckit^+^, DN1d cells were characterized as CD4^-^CD8^-^γδT^-^CD44^+^CD25^-^CD24^+^Ckit^-^, DN1e cells were characterized as CD4^-^CD8^-^γδT^-^CD44^+^CD25^-^CD24^-/low^Ckit^-^, DN2 cells were characterized as CD4^-^CD8^-^γδT^-^CD44^+^CD25^+^ and DN3 cells were characterized as CD4^-^CD8^-^γδT^-^CD44^-^CD25^+^. **D, E and F**: UMAP representation based on gene expression profiling of pooled DN1, DN2 and DN3 from WT and GFI1 KO mice colored by cluster (**D**), cell type (**E**) and genotype (**F**). **G**: Bar plots showing the distribution of each cell type found in the pool of DN1, DN2 and DN3 cells in the indicated genotypes. **H**: UMAP representation showing expression of *Maf*, *Rorc*, *Blk and Cxcr6* genes. Color represents the normalized expression of the gene. **I**: Differential gene expression showing key up-or down-regulated genes in GFI1 KO DN1-3 cells versus WT DN1-3 cells. **J**: Frequencies of c-MAF^+^RORγt^+^ cells in DN1, DN2 and DN3 cells from WT and GFI1 KO mice or control and *vav-cre GFI1^flox/flox^*mice. **K**: Representative plots showing in pink c-MAF^+^RORγt^+^DN1 cells in the DN1 subsets characterized by the expression of c-Kit and CD24 after gating on lin^-^CD4^-^CD8^-^γδT^-^CD44^+^CD25^-/low^ from WT and GFI1 KO thymus. (Lin^-^ = CD19, CD11b, CD11c, NK1.1). Adult mice meaning mice older than 5 weeks. Statistical testing was performed with multiple paired *t*-test using Prism v9 (GraphPad). *p*-Values are shown as * < 0.05, ** < 0.01, *** < 0.005, and **** < 0.001 where each statistical significance was found.

It is already known that most γδ T cells differentiate from thymic DN2 or DN3 cells^40,56^, but recently DN1d and DN1e subsets have also been shown to generate γδ T cells^40,41^. The DN2 and DN3 subsets are strongly decreased in GFI1 KO mice compared to WT animals and the DN1a/b subpopulations are almost absent (**Fig. 6C**). Even though DN1d and DN1e subsets were significantly decreased in numbers in GFI1 KO mice, their frequencies were increased in *Gfi1* deficient mice compared to control animals (**Suppl. Fig 6D**).

To gain more insight into the origin of the expanded γδT17 cell population in GFI1 KO mice, we performed a scRNA-seq analysis of pools of DN1, DN2, and DN3 cells from adult GFI1 WT and KO thymus. Our analysis showed that these cells fall into 9 distinct clusters (**Fig. 6D & Suppl. Fig. 6E & 6F**), which could be annotated as DN1, DN2 and DN3 cells and their subtypes^41,62,66^ (**Fig. 6E & Suppl. Fig. 6E**). Interestingly, clusters 6 and 8 contained populations of the DN1d and DN1e subsets, that were identified as *Rorc^+^*DN1d and *Rorc^+^*DN1e, express genes associated to the γδT17 subtype such as *Maf*, *Rorc*, *Blk, Cxcr6* and *Scart1* and were largely found in adult GFI1 KO thymus (**Fig. 6D-H**). Another cluster (cluster 5) identified as *Il2rb^+^*DN1e was also overrepresented in GFI1 KO thymus compared to WT thymus (**Fig. 6D-G**). These data show that DN1d and DN1e cells were still present in adult GFI1 KO mice, with *Rorc*^+^DN1e cells being the most expanded population in GFI1 KO DN population (**Fig. 6G**). This correlated with the high *Gfi1* expression levels found in DN1e cells compared to other DN1 cells (**Suppl. Fig. 6G**).

The most up regulated genes in the pool of DN1-DN3 cells from adult GFI1 KO thymus compared to control DN1-DN3 cells contained genes related to Vγ6^+^ γδT17 cells, such as *Maf* and *Scart1* (**Fig. 6I**), confirming that a γδT17 program was already active in DN1 cells from GFI1 KO mice. We confirmed by flow cytometry, the higher proportions of c-MAF^+^ cells in GFI1 KO and in *vav-cre Gfi1^fl/fl^* DN1e, DN2 and DN3 cells (**Suppl. 6H**) and identified an expanded c-MAF^+^RORγt^+^ DN1e population in GFI1 KO and in *vav-cre Gfi1^fl/fl^* thymus as compared to controls (**Fig. 6J & K).** This c-MAF^+^RORγt^+^ DN1e population overlapped with the abnormal DN1 population that we had described previously^17^ (**Fig. 6K**), and which expressed high level of CD44 and low level of CD25. This suggested that GFI1 restricts or controls a DN1e population bearing the potential to give rise to Vγ6^+^ γδT17 cells.

### Characterisation of the DN1 MAF^+^/RORγt^+^ and DN3 compartments in GFI1 KO mice

To determine if this DN1e subpopulation that was expanded in GFI1 KO mice can preferentially differentiate into Vγ6^+^ γδT17 cells, we performed a trajectory inference analysis using Partition-based Graph Abstraction (PAGA) by combining the scRNA-seq data obtained with DN1-3 cells and the Trgv6^+^ cluster from WT and GFI1 KO thymic γδ T cells (**Fig. 7A-C**). PAGA can infer developmental trajectories from scRNA-seq data by calculating the probability of transition between cells^67^. This provides indications about relationships and connections of cells and their precursors. Using this method, a potential trajectory was obtained from cluster 7 containing *Rorc*^+^DN1e, almost exclusively found in GFI1 KO thymus, to the three clusters containing the Vγ6^+^γδT17 cells identified as Trgv6^+^ (clusters 11, 12 and 13) that represented almost the totality of γδ T cells in GFI1 KO thymus. This indicated that the *Rorc*^+^DN1e subset enriched in GFI1 KO mice is developmentally linked to the γδT17 cells expressing the Vγ6 chain.

**Fig. 7:**
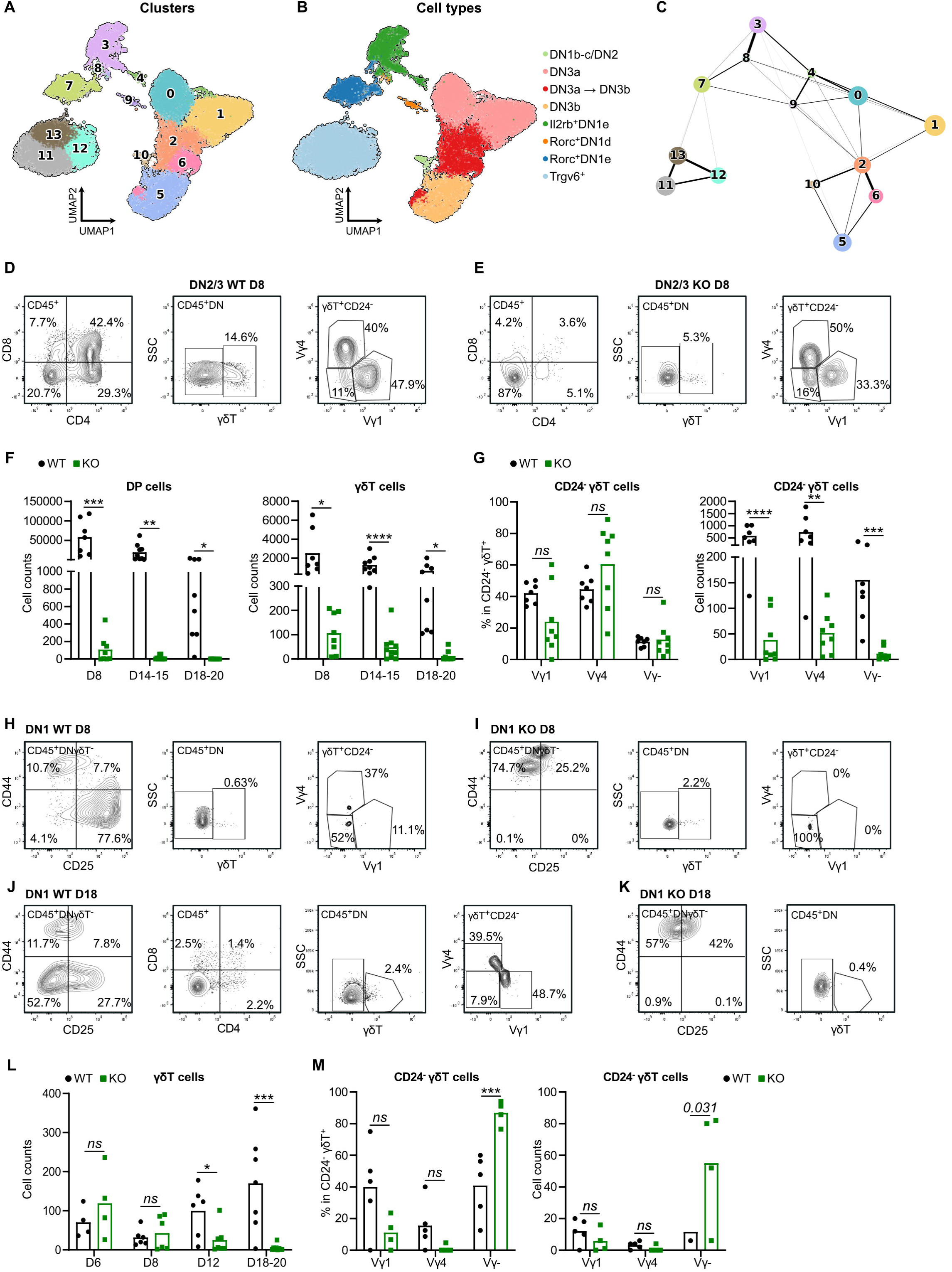
Potential of differentiation of precursors in thymus of adult mice A & B: UMAP representation based on gene expression profiling by sc-RNA-seq of thymic DN1, DN2, DN3 and γδ T cells from WT and GFI1 KO mice colored by cluster (**A**) and cell type (**B**). **C:** Partition based Graph Abstraction (PAGA) graph after running PAGA on thymic DN1-3 pre-T cells and Trgv6^+^ thymic γδ T cells from WT and KO at cluster resolution. **D** &**E**: Representative plots showing the DP (CD45^+^CD4^+^CD8^+^) population, the γδ T cells (CD45^+^CD4^-^CD8^-^γδT^+^) and the CD24^-^γδ T subsets according to the expression of Vγ chains after gating on live singlets CD45^+^CD4^-^CD8^-^γδT^+^CD24^-^ cells from sorted WT (**D**) and GFI1 KO (**E**) DN2/3 cells cultured on OP9-DL4 as described in Supp. Fig. 7I. **F**: Cell counts of generated DP cells (CD45^+^CD4^+^CD8^+^) and γδ T cells (CD45^+^CD4^-^CD8^-^γδT^+^) from WT and GFI1 KO DN2-3 cultured on OP9-DL4 as described in Supp. Fig. 7I at different time points of the coculture. **G**: Cell counts and frequencies of the indicated subsets based on the expression of Vγ chains of mature CD24^-^ γδ T cells after coculture of sorted DN2-3 cells from WT or GFI1 KO mice on OP9-DL4 cells as described in Supp. Fig. 7I. **H-K**: Representative plots showing the DN subsets based on CD44 and CD25 expression after gating on CD45^+^CD4^-^CD8^-^γδT^-^ cells, DP (CD45^+^CD4^+^CD8^+^) population, the γδ T cells (CD45^+^CD4^-^CD8^-^γδT^+^) and the CD24^-^γδ T subsets according to the expression of Vγ chains after gating on live singlets CD45^+^CD4^-^CD8^-^γδT^+^CD24^-^cells from sorted WT (**H-J**) and GFI1 KO (**I-K**) DN1 cells cultured on OP9-DL4 as described in Supp. Fig. 7 at different time points D8 (**H-I**) and D18 (**J-K**). **L**: γδ T cell counts generated from WT or GFI1 KO DN1 cultured on OP9-DL4 as described in Supp. Fig. 7 at different time points. **M**: Cell counts and frequencies of the indicated subsets of mature CD24^-^ γδ T cells after coculture of sorted DN1 cells from WT or GFI1 KO mice on OP9-DL4 cells as described in Supp. Fig. 7I. Adult mice meaning mice older than 5 weeks. Statistical testing was performed with paired *t*-test and multiple paired *t*-test using Prism v9 (GraphPad). *p*-Values are shown as * < 0.05, ** < 0.01, *** < 0.005, and **** < 0.001 where each statistical significance was found.

The cluster containing DN1-DN2 cells (cluster 10) was connected to the different DN3a and/or DN3b clusters (clusters 2 and 5) but was not linked to the Trgv6^+^ clusters (clusters 11, 12 and 13) containing Vγ6^+^γδT17 cells (**Fig. 7A-C**), reinforcing the idea that the expanded Vγ6^+^ γδT17 population in GFI1 KO thymus does not originate from the DN2 or DN3 compartment but most likely from the DN1e subset. The analysis of *Cd2-cre*; Gfi1^flox/flox^ mice, where a deletion of *Gfi1* occurs after the DN1 stage in the DN2-3 precursors during T cell development didn’t show changes in numbers for γδ T cells negative for Vγ1 and Vγ4 chain expression, expressing RORγt^+^ or producing IL17A compared to control mice (**Suppl. Fig. 7A-H**). One of the differences was decreased numbers of Vγ1^+^ γδ T cells in the thymus and in the spleen of the *Cd2-cre* Gfi1^flox/flox^ mice compared to control mice, meaning that GFI1 could have an impact on the production of this particular γδT cell subset when deleted at the DN2/3 stage (**Suppl. Fig. 7B-D**). We also observed a slightly increased proportion of RORγt^+^ cells in the spleen and a slightly increased frequency of IL17A^+^ cells in the thymus, but nothing comparable to what was found in the total GFI1 KO mice or in the *vav-cre Gfi1^fl/fl^* mice. This strongly suggests that the DN2/3 T cell precursors are not the source of the accumulated Vγ6^+^ γδT cell population seen in mice lacking GFI1. This was also confirmed when we cultured DN2/3 precursors on OP9-DL4 cells in the presence of IL7 and FLT3 (**Suppl. Fig. 7I-J**). After several days of coculture, WT DN2/DN3 cells differentiated as expected mostly into DP cells, while GFI1 KO DN2/DN3 cells were still mostly found in the DN compartment and did not differentiate into DP cells (**Fig. 7D-F**). Both WT and GFI1 KO DN2/DN3 cells were able to produce γδ T cells, but DN2/DN3 from GFI1 KO thymus produced a smaller amount of γδ T cells compared to WT DN2-3 cells (Fig. **7D-F**) reducing the numbers of the CD24^-^γδT cell subtypes (**Fig 7G**). The γδ T cells differentiated from DN2/3 lacking GFI1 did not show a preference to generate higher proportions of CD24^-^Vγ1^-^Vγ4^-^γδ T cells over the other subtypes but produced all γδ T cell populations in similar proportions to WT DN2-3 cells (**Fig. 7D, 7E & 7G**). We further tested whether a developmental link exists between GFI1 KO DN1 cells and the Vγ6^+^ γδ T cells by also culturing sorted DN1 cells from WT or GFI1 KO mice on OP9-DL4 cells in the presence of IL7 and FLT3 (**Suppl. Fig. 7I-J**). As expected, WT DN1 cells were able to progress though the different stages of T cell differentiation, while GFI1 KO DN1 remained mostly in the DN1 stage after 8 or 18 days of culture (**Fig. 7 H-K**). Only very few cells developed into γδ T cells from WT DN1 and GFI1 KO DN1 cells, with a peak of γδT cell production after 12 days for WT precursors, while GFI1 KO DN1 cells produce most of the γδ T cells earlier in the coculture before being lost (**Fig. 7L**). We gated the cells surviving in the coculture on CD24^-^γδT cells and found that the GFI1 KO did indeed produce higher proportions of Vγ1^-^Vγ4^-^ γδ T cells (most likely meaning Vγ6^+^) compared to WT DN1 cells (**Fig. 7M**). However, the number of cells produced was very low (**Fig. 7M**) suggesting that GFI1 KO DN1 cells most likely require other signals to either generate the Vγ6^+^ γδ T cells or to fully support this biased γδ T cell population leading to their accumulation in adult Gfi1 deficient mice. Our data indicate that the expanded γδT17 cell population in GFI1 KO mice are unlikely to originate from the DN2-DN3 cells. Instead DN1 subsets, particularly DN1e may contribute to this bias. However, this interpretation remains tentative, as additional experiments are required to confirm the developmental origin of these cells.

### GFI1 can directly regulate *Maf*

The scRNA-seq analysis indicted that the expression of *Maf* was increased in clusters from GFI1 KO cells and flow cytometry data confirmed this for the c-MAF protein, known to be a regulator of γδT17 cell differentiation and maintenance^68^. We also confirmed overexpression of MAF in GFI1 KO Vγ1^-^Vγ4^-^ thymic γδ T cells and also in Vγ1^+^ thymic γδ T cells compared to WT control cells (**Fig. 8A & 8B**). However, this upregulation of MAF was only associated with a higher expression or RORγt in the Vγ1^-^Vγ4^-^ thymic γδ T cells (**Fig. 8A & 8B**). We observed similar data in splenic γδ T cells, with an upregulation of c-MAF in GFI1 KO Vγ1^-^Vγ4^-^ and Vγ1^+^ splenic γδ T cells, associated with a strong RORγt upregulation only in Vγ1^-^Vγ4^-^ splenic γδ T cells (**Fig. 8C & 8D**). We also found that total DN3 cells from GFI1 KO mice that have a very different transcriptome profile compared to WT DN3 cells showed up regulation of *Maf* (**Suppl. Fig. 8A**), which was also confirmed by flow cytometry for DN2 and DN3 cells (**Fig. 8E**).

**Fig. 8:**
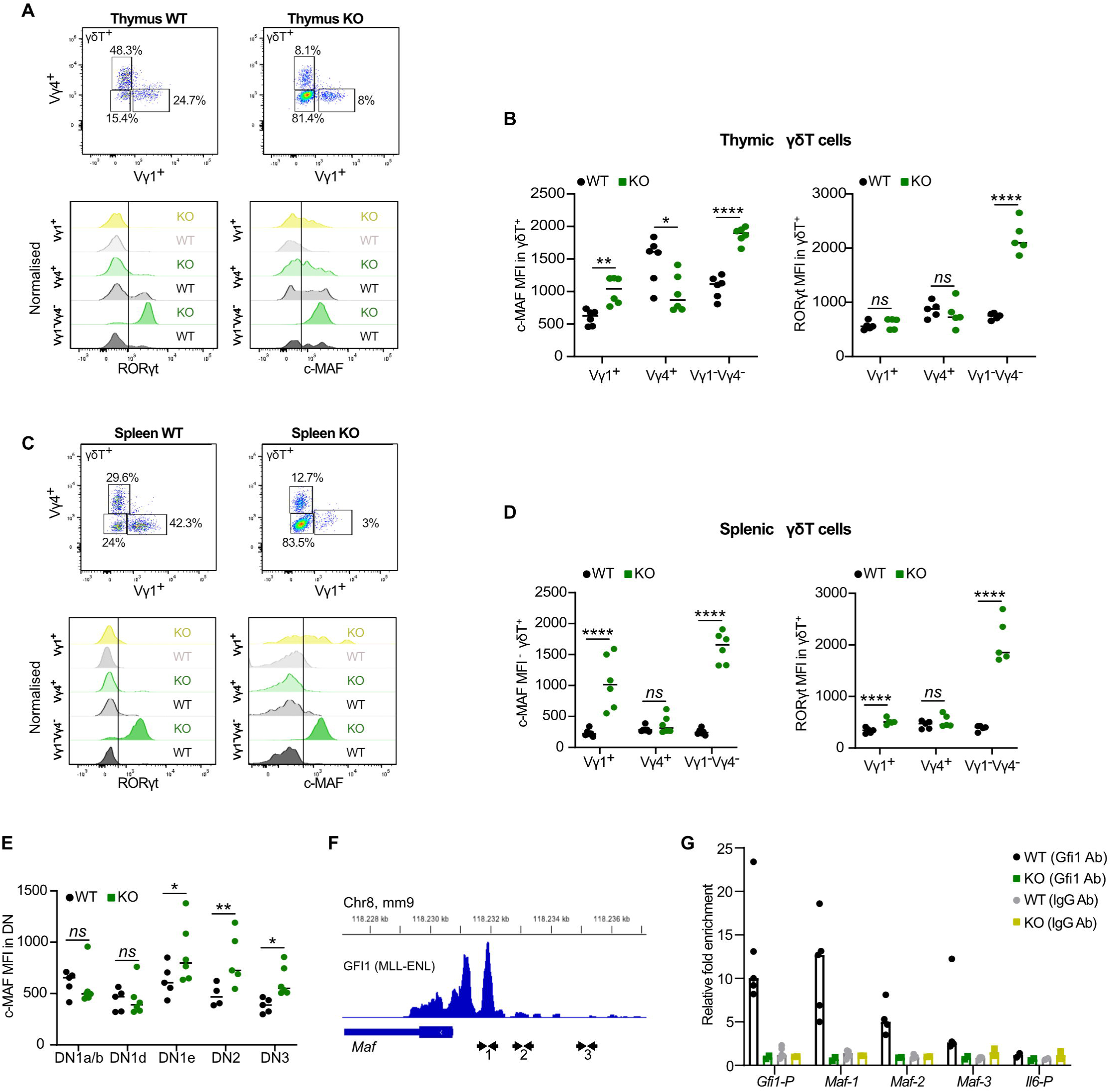
GFI1 regulation in adult mice A: RORγ and c-MAF MFI in the indicated subsets of thymic γδ T cells based on Vγ1 and Vγ4 expression from WT and GFI1 KO mice. **B**: c-MAF and RORγt MFI quantification for the indicated thymic γδ T cell subsets comparing WT and GFI1 KO mice. **C:** RORγ and c-MAF MFI in the indicated subsets of splenic γδ T cells based on Vγ1 and Vγ4 expression from WT and GFI1 KO mice. **D**: c-MAF and RORγt MFI quantification for the indicated splenic γδ T cell subsets comparing WT and GFI1 KO mice. **E**: c-MAF MFI quantification comparing WT and GFI1 KO mice for the indicated thymic DN progenitors. **F**: Peaks for GFI1 on the *Maf* promoter region from the MLL-ENL ChIP-Seq (GSE31657). **G**: ChIP-PCR on different potential binding sites (arrowheads) on *Maf* promoter for GFI1 identified in Fig. 8F with WT and GFI1 KO DN cells. *Gfi1* promoter (*Gfi1-P*) was used as positive control and *Il6 promoter* region was used as negative control. Statistical testing was performed with multiple paired *t*-test using Prism v9 (GraphPad). *p*-Values are shown as * < 0.05, ** < 0.01, *** < 0.005, and **** < 0.001 where each statistical significance was found.

In addition, we also found higher expression of c-MAF specifically in the GFI1 KO DN1e cells compared to WT cells (**Fig. 8E**), but not in DN1d or DN1a/b cells. This correlated with the high expression of GFI1 shown in DN1e cells (**Suppl. Fig. 6G)**. This data suggested that *Maf* is a potential direct GFI1 target. To test this hypothesis, we interrogated available ChIP-Seq data from MLL-ENL lymphoblastic cells and observed several peaks with a GFI1 binding motif binding in the *Maf* 5’ region and peak for GFI1 (**Fig. 8F**). Binding of GFI1 on *Maf* promoter was also found in other available ChIP-Seq data with different cell types (**Suppl. Fig. 8B**). A query of the JASPAR database confirmed the location of several GFI1 binding motifs of the *Maf* gene from the transcription start site up to several kb upstream (**Fig. 8F**). ChIP-qPCR with purified WT and GFI1 KO DN cells and primers covering these putative binding sites indicated binding of GFI1 located in regions where the primer pairs were designed (**Fig. 8G**). As controls, we used GFI1 KO DN cells, where no enrichment was found in this region and the promoter, where GFI1 does not bind, confirming occupation of the Maf promoter by GFI1 in WT DN pre-T cells (**Fig. 8G**). This suggested that GFI1 directly represses the expression of *Maf* and with this the expansion of Vγ6^+^ γδ T cells.

## Discussion

In this study, we describe expression of GFI1 in γδ T cells and their precursors and present evidence for a novel regulatory role of this transcription factor in restricting γδ T cell fate and cellularity across various tissues, including thymus, spleen, lung, and lymph nodes. Our findings demonstrate that genetic ablation of GFI1 results in a substantial expansion of RORγt^+^Vγ6^+^γδT cells producing IL17 and a concurrent decrease in Vγ1^+^ and Vγ4^+^ γδT cells. This bias toward the γδT17 subtype emerges postnatally in GFI1 KO mice and persists in the adult thymus, where the Vγ6^+^ subset secreting IL17 becomes the predominant population among γδ T cells. Furthermore, GFI1 KO mice exhibit an increased number of c-MAF^+^/RORγt^+^ cells in the DN1e subset, which has been described as containing precursors for γδT17 cells^40,41,62^. *Gfi1* deficient DN thymocytes and γδ T cells exhibit an upregulation of the bZip transcription factor c-MAF, which is known to regulate γδT17 fate^68^. We further demonstrate that GFI1 occupies the promoter region of the *Maf* gene at a cognate GFI1 binding site, suggesting that GFI1 controls the cellularity of γδ T cells and their progenitors through the potential regulation of a c-MAF-dependent transcriptional network.

The specific expansion of only Vγ6^+^ γδT cells without affecting the fate of other γδT subtypes (Vγ4^+^), which could also adopt a γδT17 profile^33,57,69^, supports the idea of an intrinsic effect of GFI1 in γδ T cell development. In addition, the experiments with *vav-cre Gfi1^fl/fl^* mice, which delete *Gfi1* only in hematopoietic cells, also ruled out that the non-hematopoietic environment such as epithelial cells in GFI1 KO mice are causing the specific expansion of Vγ6^+^ γδT17 cells seen in these animals. Transplantation experiments of WT cells into GFI1 KO animals did not reproduce the strong bias towards the γδT17 subtype observed in GFI1 KO mice, even though we could observe a slight increase of both IL17A and RORγt frequencies in donor WT γδ T cells in the spleen of the GFI1 KO recipients. This effect could be due to the transplantation itself or may suggest that the environment of the GFI1 KO spleen could have a slight impact on the fate of Vγ4^+^ γδ T, which is the other γδ T subset capable of becoming γδT17 cells.

However, it remained unclear from which cell population the expansion of Vγ6^+^ γδT17 in GFI1 KO mice originates. We could show that Vγ6^+^ γδT17 cells start to increase in numbers in GFI1 KO mice only after birth. It is thus possible that *Gfi1-*deficient precursors from the fetal thymus for Vγ6^+^ γδT17 cells continue to produce these cells when they reach adult tissues, whereas their WT counterparts would cease their production. Transplantation of adult BM cells from GFI1 KO mice did not recapitulate the expansion of Vγ6^+^ γδT17 cells in recipient animals (data not shown), which excluded the presence of a γδ T precursor within the adult thymic DN pre-T cell population. It is thus more likely that precursor cells from fetal thymus give rise to a c-MAF^+^/RORγt^+^ cells in the DN1e subset in the adult thymus where they are under the restriction of GFI1 in WT mice. These cells were seen in flow cytometry analyses in WT mice, albeit at very low numbers. It is conceivable that these cells expand into Vγ6^+^ γδT17 cells in the thymus when GFI1 is absent. Both scRNA-seq and flow cytometry show this expanded population of c-MAF^+^/RORγt^+^ cells in the DN1e pre-T cells from GFI1 KO mice very clearly. The transcription pattern of this DN1e subpopulation in GFI1 KO mice supports this notion and suggests that a DN1e subset exists with a specific transcriptional signature which is maintained after birth in the adult thymus that is associated with Vγ6^+^ γδ T cells producing IL-17. Although these data point to a fetal origin of the expanded Vγ6^+^ γδT17 cell population in GFI1 KO mice, further experimentation with fetal and adult pre-T cells are necessary to firmly support such a conclusion.

The decrease of Vγ1^+^ and Vγ4^+^ γδ T cells that we observed in GFI1 KO mice alongside the expansion of Vγ6^+^ γδT17 could be a consequence of the decreased cell numbers in the DN3 subset. Lower numbers of Vγ1^+^ and Vγ4^+^ γδ T cells were observed in fetal and adult thymus of GFI1 KO mice as compared to WT controls, in contrast to Vγ6^+^ γδ T cells, which expanded only after birth. It is thus likely that separate precursors exist for the Vγ1^+^ and Vγ4^+^ γδ T cells and the Vγ6^+^ subset. In addition, DN3 cells lacking GFI1 have a distinct transcriptomic profile from their WT counterparts, which could also affect the production of Vγ1^+^ and Vγ4^+^ γδ T cells, since they develop from this thymic DN population. Surprinsingly, an upregulation of MAF was also observed in the thymic and splenic Vγ1^+^ γδ T cells but not in the Vγ4^+^ γδ T cells, the other population where MAF can be expressed. We didn’t observe a differential expression of GFI1 (data not shown) between the different γδ T cell populations which could have explained this difference in MAF regulation. Most likely, additional regulatory or compensatory pathways exist that regulate MAF expression in different γδ T cell subtypes in addition to GFI1.

A DN1-like (CD117^−^CD24^−^CD25^−^CD44^+^) precursor subpopulation (DN1e) was more recently identified to be programmed or pre-committed to become Tγδ17 cells independently of TCR signaling^40,41^. These cells express *Rorc, Sox4, Tcf7, Tcf12, Maf, Il7r, Scart2*, and *Blk* and are predisposed to become CCR6^+^ IL17A-producing cells^41^. c-MAF promotes the expression of genes essential for Tγδ17 cell identity, such as *Ror*γ*t* and *Blk*, which are key players in the Tγδ17 program, and which are also found to be increased in expression in GFI1 KO γδT cells^68,70^. Our data suggest that lineage committed precursors and γδT17 cells are not only controlled by TCF1 and HEB/SOX13^71,72^, which regulate c-MAF, but also by GFI1, which acts upstream of c-MAF. This would be coherent with the model that lineage fate can be prewired cell-intrinsically and is not necessarily specified by antigen receptor signals^41^. This also lends further support to the view that the MAF^+^/ RORγt^+^ cell population that we found expanded in thymic DN1e cells from GFI1 KO mice represents the precursor for the expanded IL17 producing Vγ6^+^ γδ T cells. However, further experimentations would be needed to support this hypothesis, because the coculture of DN1 cells from GFI1 KO mice on OP9-DL4 layer or in FTOC (data not shown) was not robust enough to support the expansion of Vγ6^+^ γδ T, even though a bias was observed towards this subtype. It may suggest that other signals in the periphery such as the cytokine environment specifically found in GFI1 KO mice may be required to support the intrinsically biased γδ T populations to expand in *Gfi1* deficient mice.

It has previously been shown that c-MAF is highly expressed in RORγt^+^ γδ T cells, which are γδT17 cells, but not in other γδ T cell subsets like γδT1 cells^68^. c-MAF functions by promoting the expression of key genes in the type 17 program (e.g., *Rorc, Il17a, Blk*) and possibly also *Il23R* and antagonizes negative regulators like TCF1, which are required to upregulate *Ifng*. c-MAF helps γδ T cells adapt to specific tissue microenvironments by regulating metabolic pathways and integrates signals from IL1β, IL23 and interactions with epithelial cells to fine-tune γδ T cell responses^73^. These reports identified c-MAF as a universal and essential regulator of γδT17 cell differentiation and function but still lacked insights into the molecular mechanisms that control c-MAF expression.

Our results with ChIP-qPCR and ChIP-seq data from the public domain showed that GFI1 occupies the *Maf* promoter region. This observation together with our findings that c-MAF expression is upregulated in all tested *Gfi1* deficient γδ T cells indicates that GFI1 acts as an upstream negative regulator of c-MAF. It is therefore conceivable that the direct transcriptional repression of the *Maf* gene by GFI1 is necessary to maintain and restrict the development of Vγ6 γδT17 cells and that absence GFI1 leads to the developmental defect in these γδ T cells seen in GFI1 KO animals. The upregulation of c-MAF and c-MAF target genes such as *Il17, Blk* support such a role of GFI1, but further studies are needed to establish a direct mechanistic link.

It is also known that GFI1 antagonizes RORγt expression through GATA3^3,23,70^ and that-c-MAF induces expression of RORγt and synergizes with RORγt to regulate IL17-producing cells. It is thus conceivable that a GFI1/c-MAF/RORγt regulatory circuit exists that controls the expression of IL17A and ensures the cellularity and functionality of γδT17 cells. GFI1 has been shown to regulate Notch/IL7 signaling^18^, both important for γδ T cells, suggesting that it could be involved at different levels, possibly also in γδ T cell development. Our data do not exclude contribution from these pathways and further experiments would be required to decide whether the link between Notch/IL7 signaling and GFI1 bears any role in γδ T cells.

The ability of c-MAF to fine-tune IL17 production has been highlighted as a potential therapeutic target for controlling γδ T cell-mediated inflammation without compromising their protective roles^74^. Strategies to modulate c-MAF activity or its interaction with RORγt are being explored as therapeutic options to balance inflammation and immunity. Our findings which place GFI1 upstream of c-MAF may point to new possibilities to interfere with this regulatory circuit since GFI1 is in a complex with LSD1 for which small molecule inhibitors exist^75^.

## Materials and Methods

### Mice

The generation of GFI1 KO, GFI1-EGFP, and GFI1-fl/fl mice was previously described^13,28,63^. C57BL6/J CD45.2 congenic mice (B6.SJL-*Ptprc^a^ Pepc^b^*/BoyJ) and *Vav-cre* mice (B6.Cg-Tg(VAV1-cre)1Graf/MdfJ) were obtained from Jackson labs (Bar Harbour, MN). C57BL6/J CD45.1/2 mice were a gift from Dr. Javier Di Noia, IRCM, Montreal and hCD2-iCre mice (B6.Cg-Tg(CD2-icre)4Kio/J) were a gift from Dr. André Veillette, IRCM, Montreal. All mice were maintained on the C57BL6/J background. Mice were housed and bred in the IRCM specific pathogens-free facility in a 12 h dark / light cycle with food and drink at libitum. All mouse work was reviewed and approved by the animal protection committee at the IRCM (protocols #2020-09), according to the guidelines of the Canadian Council of Animal Care.

### Tissue dissociation

Lungs from adult mice (at least 5 weeks old) were perfused with 10 ml of PBS *via* the right ventricle and then were cut into small pieces with scissors and a scalpel. Lung pieces were dissociated and digested in a buffer with collagenase IV (1 mg/ml) (Gibco) and DNase I (0.2 mg/ml) (Sigma) in RPMI medium for 30 min at 37°C. Cells were aspirated several times with a syringe and needle (18 gauge) every 10 min to further the dissociation and digestion process. Lung cell suspensions were filtered and washed in PBS. Thymi, spleen, and lymph nodes from adult mice (at least 5 weeks old) were mechanically dissociated to obtain a cell suspension which were filtered. Then, red blood cell lysis buffer (Sigma) was added to the lung, thymic, splenic or LN cell pellet for 4 min and washed before staining or sorting.

### Bone marrow chimera

Bone marrow (BM) cells from CD45.1/2 mice were obtained by flushing femurs and tibiae with cold PBS using a syringe with 23G needle. The red blood cells were subsequently lysed, and the cells were washed in PBS before being injected by the tail vein in 100 µl of PBS containing 2×10^6^ BM cells into either CD45.2 WT or GFI1 KO recipient mice. 4 or 5 weeks after the transplantation, the recipient mice were sacrificed, and splenic, thymic, and lung cells were analyzed by flow cytometry.

### OP9-DL1 and OP9-DL4 culture

OP9-DL1 and OP9-DL4 cells were a gift from the lab of Dr. Zúñiga-Pflücker^76^. OP9-DL1 cells were plated in 96 flat-bottom well plate at a density of 3000 cells in 100 uL of OP9 medium (AMEM, 20% FBS, 1% Pen/strep, 2.2g/L sodium bicarbonate, 50 μM β-mercaptoethanol). The day after, the same numbers of WT and GFI1 KO sorted DN1, DN2-DN3 or total DN cells were co-cultured on OP9-DL1 or OP9-DL4 monolayers in DMEM-glutamax medium containing 10% of FBS, 1% Pen/strep, 50 μM β-mercaptoethanol, 10 mM Hepes with Flt3 (5 ng/mL, Peprotech) and IL7 (1 ng/mL, Peprotech). Thymocytes were transferred to a new OP9-DL1 or OP9-DL4 monolayer with fresh cytokines every 3 days.

### Flow cytometry and cell sorting

For the analysis of cells from spleen, lung, lymph nodes and thymus from mice by flow cytometry, a Fortessa (BD Biosciences) flow cytometer and FlowJo software were used. Cell sorting was performed on a FACSAria III Sorter (BD Biosciences). For IL17A and IFNγ intracellular staining, cells were cultured for at least 3h with GolgiPlug (BD Biosciences), PMA (50μM, Sigma) and ionomycin (1mM, Sigma) and stained using the eBioscience cytokine intracellular staining kit (Invitrogen). For IL22 staining^77^, cells were first cultured with IL23 (20 ng/mL, R&D Systems) overnight before stimulation with PMA (50μM, Sigma) and ionomycin (1mM, Sigma) in the presence of GolgiPlug (BD Biosciences) for the last 4 hours. Intracellular staining for the transcription factors RORγt and c-MAF was also done using eBioscience FoxP3 intracellular kit (Invitrogen). For phosphoSTAT3 intracellular staining, after the surface staining, cells were stimulated for 15 minutes at 37°C with either IL23 (20ng/mL, R&D Systems) or IL7 (10 ng/mL, Peprotech) and washed with cold PBS. Cells were subsequently fixed for 15 minutes at 37°C with Biolegend fixation buffer^78^ before being permeabilized with the Biolegend True-phos Perm buffer (425401) for 30 minutes at 4°C. After washing, the cells were incubated with Phospho-STAT3 antibody for 30 min at 4°C. For the splenic γδ T cell culture, sorted splenic γδ T cells were cultured *in vitro* in DMEM medium containing 10% FBS, 1% Pen/Strep, 10 mM Hepes and 50μM β-mercaptoethanol in the presence or not of IL23 (20 ng/mL, R&D Systems) and IL1β (10 ng/mL, Biolegend) for 16 hours and GolgiPlug (BD Biosciences) was added for the last 4 hours, before cytokine staining. All antibodies directly conjugated to a fluorochrome or to biotin were purchased from BioLegend, eBiosciences, or BD Biosciences (Table 1).

### PCR

RNA extraction was performed using Tri-Reagent (Life Technologies) for cell numbers between 1 × 10^6^ and 3 × 10^6^ or using the RNeasy Micro kit from QIAGEN for low numbers of cells following the company’s instructions (RNeasy Micro kit; QIAGEN). RT-PCR was performed using Superscript II (Invitrogen). RNA was incubated with oligodeoxy thymidine (Invitrogen), EDTA (Invitrogen), and dNTP (Invitrogen) for 10 min at 65°C and then DTT (Invitrogen), Superscript buffer 5× (Invitrogen), RNaseOUT buffer (Invitrogen), and the enzyme Superscript II (Invitrogen) were added and the reaction was subjected to PCR cycles (step 1, 25°C for 10 min; step 2, 42°C for 60 min; step 3, 70°C for 15 min) to generate cDNA for real-time quantitative PCR. Real-time quantitative PCR was performed in triplicates on the ViiA7 real-time PCR machine (Applied Biosystems) in SYBR Green Master Mix (Applied Biosystems) containing specific primers for mouse. The primers used are *Gfi1 Forward*: GTGCCATCAGAGGAGGTGAA, *Gfi1 Reverse*: TCTGATAACCTGCGGCCAAT; *Gfi1b Forward*: CAGTCCTACGCCCACCTTGG, *Gfi1b Reverse*: AGTGTCCGAGTGGATGAGCA, and *Gapdh Forward*: ACTGAGCAAGAGAGGCC, *Gapdh Reverse*: TATGGGGGTCTGGGATGGAA.

### Single cell RNA-seq sequencing

The single-cell RNA sequencing on lungs was performed using 10x Genomics platform with the Chromium Next GEM Single Cell 3 Kit v3.1. Total lung cells from adult mice (between 5 and 8 weeks old) were sorted on live cells using Hoecsht to stain the dead cells. Then, cells were loaded on a Chromium controller on a chip G to target a recovery of 10,000 cells per sample and library were constructed following the manufacturer recommendations. Library size distribution was assessed on a 2100 bioanalyzer (Agilent Technologies) using a High Sensitivity DNA Kit and libraries were quantified by qPCR. Equimolar libraries were sequenced in paired-end reads (PE 28-91), on a Novaseq 6000 system (Illumina), with an average depth of 25,000 reads per cell. The single-cell RNA sequencing on sorted γδ T cells (from the spleen or the thymus DN1-DN2 of adult mice) or on sorted DN1-DN3 cells was performed using 10x Genomics platform with the Chromium Next GEM Single Cell 3 v3.1 Kit and the 3’ Feature barcode kit for cell surface protein. To accelerate the sorting step, splenic cells or thymic cells were preselected to remove B cells (CD19), myeloid cells (Gr1, CD11b and CD11c) and NK cells (NK1.1 and CD49b) using biotinylated antibodies and Mojo magnet selection (Biolegend), before FACS staining. CD8^+^ cells were also partially removed from the thymus using Mojo Magnet selection (Biolegend). Then, WT or GFI1 KO cells were stained with TotalSeq-B oligo-conjugated antibodies (Totalseq-B0302 & Totalseq-B0303, Biolegend) to hashtag the samples in order to reduce costs and decrease technical variability during the incubation with the fluorochrome coupled antibodies for the FACS sorting. Sorted cells were loaded on a Chromium controller on a chip G to target a recovery of 20,000 cells per sample (10,000 cells per condition WT or KO) and library were constructed following the manufacturer recommendations. Library size distribution was assessed on a 2100 bioanalyzer (Agilent Technologies) using a High Sensitivity DNA Kit and libraries were quantified by qPCR. Equimolar libraries were sequenced in paired-end reads (PE 100), on a Novaseq X system (Illumina). An average depth of 25,000 or 5,000 reads per cell was targeted for gene expression or hashtag libraries, respectively. Duplicatat were used for all the single cell RNA-sequencing experiments, meaning 2 WT and 2 GFI1 KO samples for each cell types.

### Single cell RNA-seq analyses

scRNAseq data was analyzed using python (v3.12.4) and the package scanpy (v1.10.4). Cells with <500 reads, <200 genes or>20% mitochondrial genes were removed as outliers. Importantly, ribosomal genes, predicted genes with *Gm-*identifier and mitochondrial genes were excluded for subsequent analysis. The samples were demultiplexed using hashsolo^79^. The datasets were normalized using the normalize_total function with a target sum of 10000, followed by log1p transformation and scaling with a maximum value of 10. The top 2000 highly variable features were calculated using the flavor “seurat_v3” and the raw counts. Dimensionality reduction was performed with default settings, while the resolution for clustering using the Leiden algorithm was set to 0.5 for the DN dataset, 0.8 for the thymic γδ T cells and lung datasets and 1 for the splenic γδ T cells dataset. To characterize the clusters, differential gene expression analysis was done using the rank_genes_groups function. Graph abstraction was performed using the function paga in scanpy with default parameters using clusters as partitions^80^.

### Chromatin immunoprecipitation (Ch-IP)

Ch-IP for GFI1 was performed on 10 × 10^6^ fresh cells. Briefly, cells were cross-linked with 1% formaldehyde for 10 min before quenching with 125mM glycine for 5 min. Cells were initially lysed in cell lysis buffer (5mM PIPES, 85mM KCl, 0.5% Nonidet P-40, freshly added protease inhibitors mixture, and 0.8mM PMSF) for 15 min on ice with occasional vortexing. After centrifugation at 4°C at 4000 rpm for 5 min, the pellet was resuspended in nuclei lysis buffer (50mM Tris [pH 8], 10mM EDTA, 0.1% SDS, freshly added protease inhibitors mixture, and 0.8mM PMSF) and incubated on ice for 20 min with occasional vortexing. The lysed cells were sonicated using a Covaris E220 to generate 200–600-bp fragments. One volume of immunoprecipitation dilution buffer (300mM NaCl and 2% Triton X-100) was added to sonicated nuclei lysates and was immune-precipitated with 5μg of anti-Gfi1 (AF3540; R&D Systems). Beads were added to harvest protein DNA complexes. After de-crosslinking, the eluate was digested with RNase A and proteinase K. The chromatin associated DNA was extracted using QIAquick PCR Purification Kit (Qiagen). The primers for the ChIP-PCR are the following: *Gfi1 Promoter Forward*: AATTCGGGAGCTGAAGGCAA, *Gfi1 Promoter Reverse*: AGAGGGGCACACAGTTTAGC; *Maf-1 Forward*: CTCTCGAACTCCGAGGAAAGC, *Maf-1 Reverse*: GAGCGGCAGCGATTAGCAT; *Maf-2 Forward*: TGGTCAATAATGACAAGCGCA, *Maf-2 Reverse*: ATACAAATCGGAACCCCGGC; *Maf-3 Forward*: TCAATCAGTGTAGAAAACTGAACC, *Maf-3 Reverse*: ACAAGTCCGGAAACCCAGAT;; *Il6 Promoter Forward:* GGCGGAGCTATTGAGACTGT, *Il6 Promoter Reverse:* AAACCGGCAAGTGAGCAGAT and *Chr2 Forward*: TGGGCATATCCCTGGAGCTT, *Chr2 Reverse*: GGCCATCCCACAGTCACAAC.

## Statistical analysis

Multiple paired *t*-test or paired *t*-test was used to calculate p values where indicated. A p value ≤0.05 was considered as statistically significant. Survival curves were analyzed by log-rank Mantel–Cox test using GraphPad Prism (GraphPad Software, La Jolla, CA). The p values are as follows: *p < 0.05, **p < 0.01, ***p < 0.005, and ****p < 0.001.

## Data availability and resources

The primary read files and the raw counts for all single-cell sequencing datasets reported in this paper are available to download from the Gene Expression Omnibus under accession number GSE294462. The analysis of sc-RNA-seq data and codes for each Figure containing sc-RNA-seq data are also deposited on GEO. Codes are available upon request. All data supporting the findings of this study are available within the paper and its Supplementary Information. Antibodies are listed on a separate table under supplementary information.

## Authors’ Contributions

J. Fraszczak: Conceptualization, data curation, formal analysis, validation, investigation, visualization, methodology, writing original draft, revising, and editing. D. Obwegs: Single cell RNA-seq analysis and interpretation. K. Arman: ChIP-PCR experiments and analysis. T. Muralt: data curation, formal analysis. Sagar: Single cell RNA-seq analysis and interpretation, writing, and editing. I King: support with materials, resources, and reading manuscript. B. Thurairajah: support with materials, resources. H. Melichar: support with reading manuscript. E. Mallet Gauthier: support with experiments. T. Moroy: Conceptualization, resources, data curation, supervision, funding acquisition, investigation, visualization, writing original draft, project administration, and editing.

## Acknowledgments

The authors thank the IRCM animal technicians, the IRCM flow cytometry facility staff for help with cell sorting, the IRCM molecular genomics facility staff for performing single cell RNA-seq experiment and advice, the IRCM bioinformatic facility for processing single cell RNA-seq data. The authors thank Mathieu Lapointe for the mouse genotyping and all the Dr Moroy’s lab members for advice. This work was supported by the Canadian Institutes for Health Research (CIHR) through a Foundation grant (FDN - 148372) and by the IRCM Foundation.

## Supplementary data

**Sup. Fig. 1: Phenotype of** γδ **T cells in adult GFI1 KO mice**

**A:** Violin plots showing the number of reads and genes as well as the percentage of mitochondrial genes quantified per cell in the lung tissue in the scRNA-seq data. **B:** Dot plot showing the key differentially expressed genes in each cluster shown in Fig. 1A. Color represents the scaled mean expression of the gene in the respective cluster, and dot size represents the fraction of cells in the cluster expressing the gene. **C**: Representative plots on small intestine (SI) and colon lamina propria cells showing γδ T cells and CD4 T cells after gating on live (according to FSC/SSC) singlets CD45^+^CD3^+^ cells. SI and colon lamina propria cells were isolated following published protocols^81^. **D** & **E**: CD4^+^ T cell counts in the spleen, LN, and lung of WT and GFI1 KO mice (**D**) or control and *vav-cre GFI1^flox/flox^* mice (**E**). Adult mice meaning mice older than 5 weeks. Statistical testing was performed with paired *t*-test using Prism v9 (GraphPad). *p*-Values are shown as * < 0.05, ** < 0.01, *** < 0.005, and **** < 0.001 where each statistical significance was found.

**Sup. Fig. 2: Expansion of the ROR**γ**t^+^IL17a^+^** γδ**T subset in adult GFI1 KO mice.**

**A**: Representative plots showing the expression of CD27 on gated live singlets CD45^+^γδT^+^CD4^-^cells from WT and GFI1 KO spleen, LN and lung. **B**: Representative normalized histograms of RORγt^+^ cells in either singlets CD45^+^CD4^-^γδT^+^ or CD45^+^CD4^+^γδT^-^ splenic, LN or lung cells from control and *Vav-cre Gfi1^flox/flox^* mice. **C**: Frequencies of RORγt^+^ cells within singlets CD45^+^CD4^-^ γδT^+^ cells from splenic, LN and lung compared to control and *vav-cre GFI1^flox/flox^* mice. **D**: splenic, LN and lung CD45^+^CD4^-^γδT^+^RORγt^+^ cell counts in control and *vav-cre GFI1^flox/flox^* mice. **E**: Representative plots showing the expression of IL22 and IL17A in gated singlets CD45^+^CD4^-^γδT^+^ cells in the spleen, LN and lung from WT and GFI1 KO mice. Cells were stimulated with IL23 (20ng/mL) overnight and treated with PMA, ionomycin and Golgi Plug for the last 4 hours before staining. **F**: Percentages of IL17A^+^IL22^+^ cells in γδ T cells from the spleen, LN and lung of WT and GFI1 KO mice. **G**: IL17A^+^IL22^+^γδT^+^ cell counts in the spleen, LN and lung of WT and GFI1 KO mice. Adult mice meaning mice older than 5 weeks. Statistical testing was performed with paired *t*-test using Prism v9 (GraphPad). *p*-Values are shown as * < 0.05, ** < 0.01, *** < 0.005 and **** < 0.001 where each statistical significance was found.

**Sup. Fig. 3: Expansion of V**γ**6^+^** γδ**T17 cells in the periphery of adult GFI1 KO mice.**

**A:** Violin plots showing the number of reads and genes as well as the percentage of mitochondrial genes quantified per cell in the splenic γδ T cells in the scRNA-seq data. **B**: Representative normalized histogram of PhosphoSTAT3 in gated WT and GFI1 KO Vγ1^-^Vγ4^-^Vγ5^-^ γδ T stimulated or not with IL7 (10 ng/mL) or IL23 (20 ng/mL) for 15 min and the measured mean fluorescence (MFI) for each condition. Adult mice meaning mice older than 5 weeks. Statistical testing was performed with multiple paired *t*-test using Prism v9 (GraphPad). *p*-Values are shown as * < 0.05, ** < 0.01, *** < 0.005, and **** < 0.001 where each statistical significance was found.

**Sup. Fig. 4: Expansion of V**γ**6^+^** γδ**T17 cells in the thymus of adult GFI1 KO mice.**

**A**: Cell numbers (left) and frequencies in CD45^+^ cells (right) of γδ T cells (CD4^-^γδT^+^) in the thymus of control and *vav-cre GFI1^flox/flox^* mice. **B**: Representative normalized histogram of RORγt^+^ cells in either CD45^+^CD4^-^γδT^+^ or CD45^+^CD4^+^γδT^-^ total thymocytes from control and *vav-cre, Gfi1*^flox/flox^ mice. **C**: Frequencies of RORγt^+^ cells in thymic CD45^+^CD4^-^γδT^+^ comparing control and *vav-cre, Gfi1*^flox/flox^ mice (left) and thymic CD45^+^CD4^-^γδT^+^RORγt^+^ cell counts in control and *vav-cre, Gfi1*^flox/flox^ mice (right). **D:** Violin plots showing the number of reads and genes as well as the percentage of mitochondrial genes quantified per cell in the thymic γδ T cells in the scRNA-seq data. **E**: Differential gene expression showing key up-or down-regulated genes in GFI1 KO thymic γδ T cells versus WT splenic γδ T cells. **F**: Representative plots showing CD44 and CD45RB expression in gated thymic CD45^+^CD4^-^γδT^+^CD24^-^ cells from WT and GFI1KO mice. **G**: Cell counts of thymic CD45^+^CD4^-^γδT^-^CD24^-^CD44^+^CD45RB^-^ (γδT17), CD45^+^CD4^-^γδT^-^CD24^-^CD44^+^CD45RB^+^ and CD45^+^CD4^-^γδT^-^CD24^-^CD44^-^CD45RB^+^ cells in WT and GFI1 KO mice. **H**: Representative plots of CD24 expression in γδ T cells and Vγ chains expression in gated CD24^-^ γδ T cells from D8-10 and D18-D20 old WT and GFI1 KO mice. **I**: Cell numbers of thymic CD24^-^ γδT^+^ cells in D8-10 and D18-D20 old WT and GFI1 KO mice. **J & K**: Vγ chain frequencies (**J**) and Vγ chain cell counts (**K**) in thymic CD24^-^ γδ T cells from D8-10 and D18-D20 old WT and GFI1 KO mice. Vγ^-^ = Vγ1^-^Vγ4^-^Vγ5^-^. Statistical testing was performed with multiple paired *t*-test for **G**, multiple unpaired *t*-test for **I**, **J** and **K** or paired *t*-test for **A** and **C**, using Prism v9 (GraphPad). *p*-Values are shown as * < 0.05, ** < 0.01, *** < 0.005, and **** < 0.001 where each statistical significance was found.

**Sup. Fig. 5: GFI1 intrinsically controls the generation of γδ T cells.**

**A**: *Gfi1b* mRNA expression in γδ T cells from thymus, spleen, LN, and lung. *Gapdh* was used as the housekeeping gene. **B**: Representative plots showing CD45.1/2^+^ donor γδ T cells and CD45.2^+^ recipient γδ T cells in the spleen, thymus and lung of non-irradiated WT and KO recipients after gating on live, singlets cells. **C and D**: Representative plots showing IL17A^+^ (C) and RORγt^+^ (D) cells previously gated on CD45.1/2^+^ donor γδ T cells or CD45.2^+^ recipient (WT and KO) γδ T cells from the spleen, thymus and lung of non-irradiated WT and KO recipients after gating on live, singlets cells. **E**: Representative plots showing Vγ1 and Vγ4 expression in CD45.1/2^+^ donor γδ T cells or in CD45.2^+^ recipient (WT and KO) γδ T cells from the spleen, thymus and lung of non-irradiated WT and KO recipients after gating on live, singlets cells.

**Sup. Fig. 6: Precursors in thymus of adult mice**

**A**: Scheme showing the culture of sorted thymic DN cells from WT and KO mice on an OP9-DL1 layer in the presence of Flt3 (5 ng/mL) and IL7 (1 ng/mL) for 12-14 days. **B**: Representative plots showing the purity of the thymic DN population before selection (total thymocytes) and after selection. **C**: Representative plots showing the purity of the thymic DN population after selection from WT and GFI1 KO thymi. **D**: DN cell frequencies in the thymus of WT and GFI1 KO mice or control and *vav-cre GFI1^flox/flox^* mice. DN1a/b cells were characterized as CD4^-^CD8^-^γδT^-^CD44^+^CD25^-^CD24^+^Ckit^+^, DN1d cells were characterized as CD4^-^CD8^-^γδT^-^CD44^+^CD25^-^CD24^+^Ckit^-^, DN1e cells were characterized as CD4^-^CD8^-^γδT^-^CD44^+^CD25^-^CD24^-/low^Ckit^-^, DN2 cells were characterized as CD4^-^CD8^-^γδT^-^CD44^+^CD25^+^ and DN3 cells were characterized as CD4^-^CD8^-^γδT^-^CD44^-^CD25^+^. **E**: Dot plot showing the key differentially expressed genes in each cluster shown in Fig. 6C. Color represents the mean expression of the gene in the respective cluster, and dot size represents the fraction of cells in the cluster expressing the gene. **F**: Violin plots showing the number of reads and genes as well as the percentage of mitochondrial genes quantified per cell in the thymic DN cells in the scRNA-seq data. **G**: GFP^+^ cells indicating Gfi1 expression in the indicated thymic DN populations from GFI1:GFP reporter mice (het) and controls (wt). **H**: c-MAF % in DN1a/b, DN1d, DN1e, DN2 and DN3 cells for WT and GFI1 KO mice or control and *vav-cre GFI1^flox/flox^* mice analyzed by flow cytometry. Statistical testing was performed with multiple paired *t*-test **G** using Prism v9 (GraphPad). *p*-Values are shown as * < 0.05, ** < 0.01, *** < 0.005 and **** < 0.001 where each statistical significance was found.

**Sup. Fig. 7: Potential of differentiation of precursors in thymus in adult mice**

**A**: Representative plots of splenocytes and thymocytes from control and *Cd2-cre* Gfi1^flox/flox^ mice analyzed by flow cytometry and stained to identify γδ T cells (CD45^+^CD4^-^γδT^+^) after gating on live cells (according to the FSC/SSC) and singlets. **B**: Cell numbers and frequencies in CD45^+^ cells of γδ T cells (CD4^-^γδT^+^) in the spleen and in the thymus of control and *Cd2-cre* Gfi1^flox/flox^ mice. **C & D**: Cell counts of Vγ1^+^ γδ T cells, Vγ4^+^ γδ T cells and Vγ1^-^Vγ4^-^ γδ T cells in the spleen (**C**) and in the thymus (**D**) of control and *Cd2-cre* Gfi1^flox/flox^ mice. **E:** Representative plots of splenic and thymic cells from control and *Cd2-cre* Gfi1^flox/flox^ mice treated with PMA, ionomycin and Golgi Plug for 3.5 hours and analysed by flow cytometry. The cells were gated on singlets CD45^+^CD4^-^γδT^+^ before gating for IFNγ and IL17A. **F:** Representative normalized histogram of RORγt^+^ cells in either splenic or thymic CD45^+^CD4^-^γδT^+^ and CD45^+^CD4^+^γδT^-^ from control and *Cd2-cre* Gfi1^flox/flox^ mice. **G**: Frequencies of IL17A^+^ cells in γδT^+^ cells and CD45^+^CD4^-^γδT^+^IL17A^+^ cell counts in the spleen and thymus of control and *Cd2-cre* Gfi1^flox/flox^ mice. **H**: Frequencies of RORγt^+^ cells in γδT^+^ cells and CD45^+^CD4^-^γδT^+^RORγt^+^ cell counts in the spleen and thymus of control and *Cd2-cre* Gfi1^flox/flox^ mice. **I**: Scheme showing the culture of sorted thymic DN1 or DN2/3 cells from WT and KO mice on an OP9-DL4 layer in the presence of Flt3 (5 ng/mL) and IL7 (1 ng/mL) for maximum 20 days. **J**: Representative plots showing the gating strategy to sort DN1 and DN2/DN3 for the coculture with the OP9-DL4 cells described in Suppl. Fig. 7I. Statistical testing was performed with paired *t*-test **G** using Prism v9 (GraphPad). *p*-Values are shown as * < 0.05, ** < 0.01, *** < 0.005 and **** < 0.001 where each statistical significance was found

**Sup. Fig. 8: GFI1 regulation in adult mice**

**A:** Differential gene expression showing key up-or down-regulated genes in GFI1 KO DN3a, DN3a→b and DN3b cells versus WT cells. **B**: Peak for GFI1 on *Maf* promoter from ChIP-seq analysis from Mast cells (GSE42518), ILC2 cells (GSE50806) and Th1 cells (GSE70893).

