## Supplementary figures and images for "GFI1 as a novel regulator of γδ T cell development and the IL-17/IFNγ lineage commitment"

### Suppl. Fig. 1

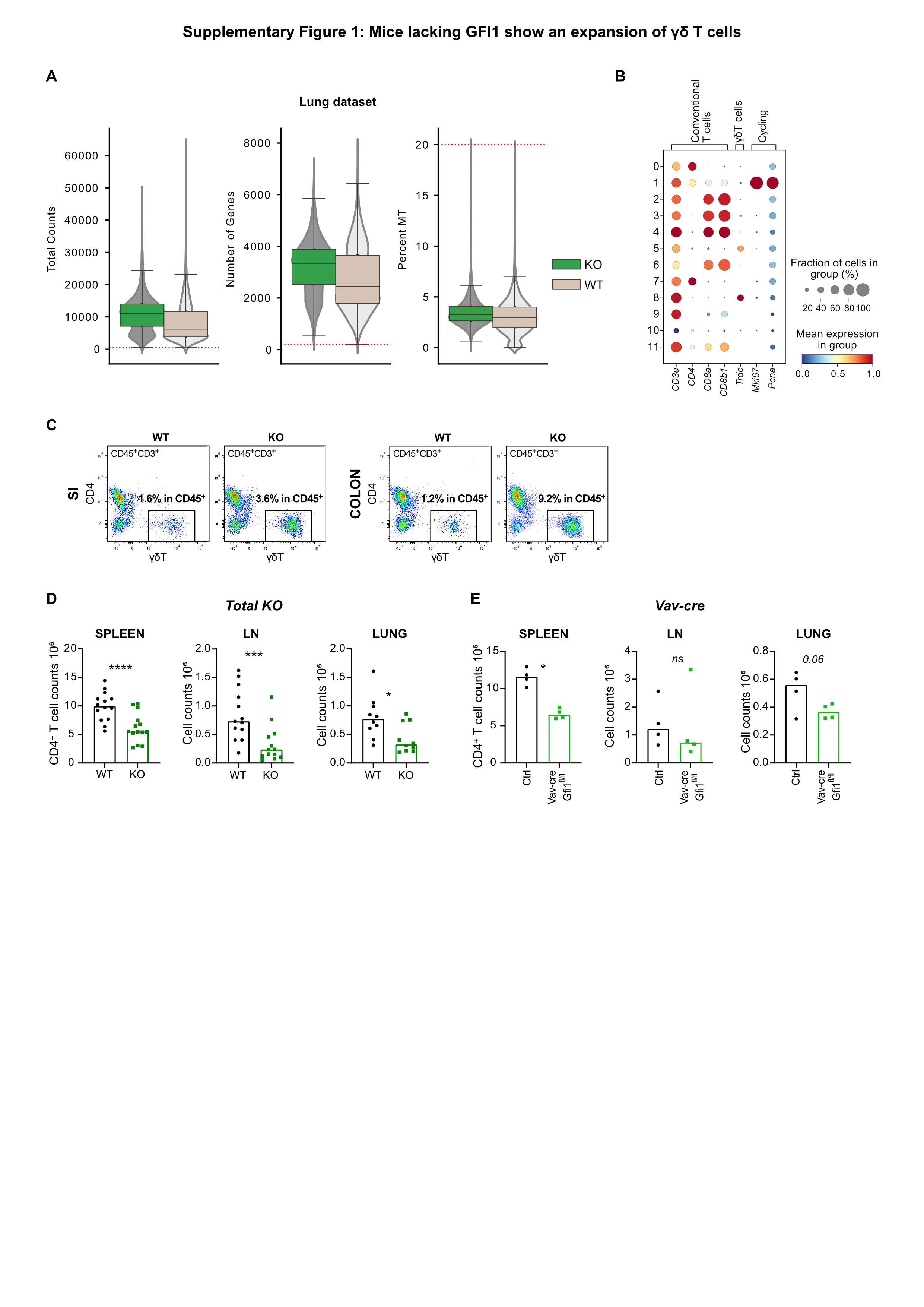

### Suppl. Fig. 2

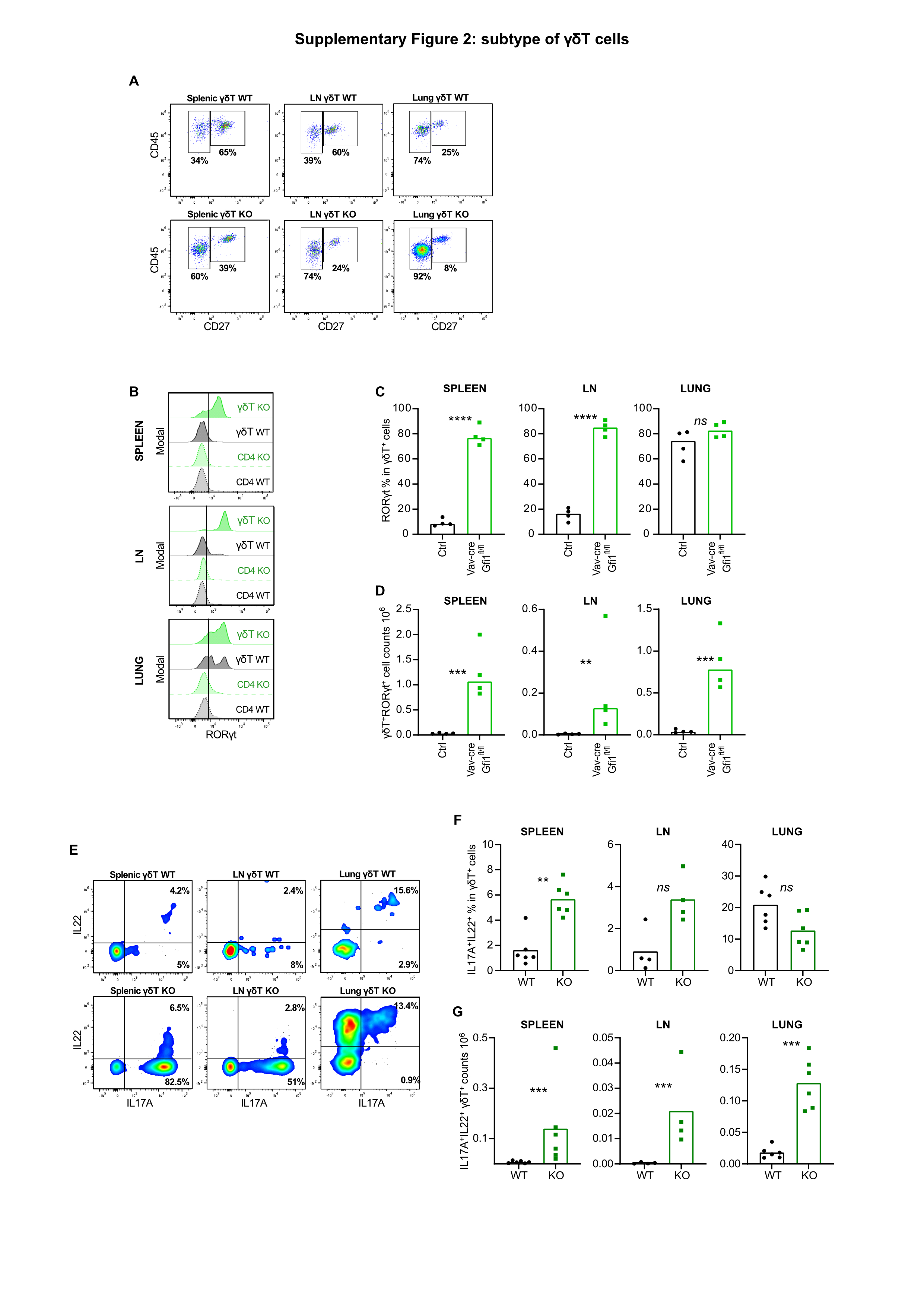

### Suppl. Fig. 3

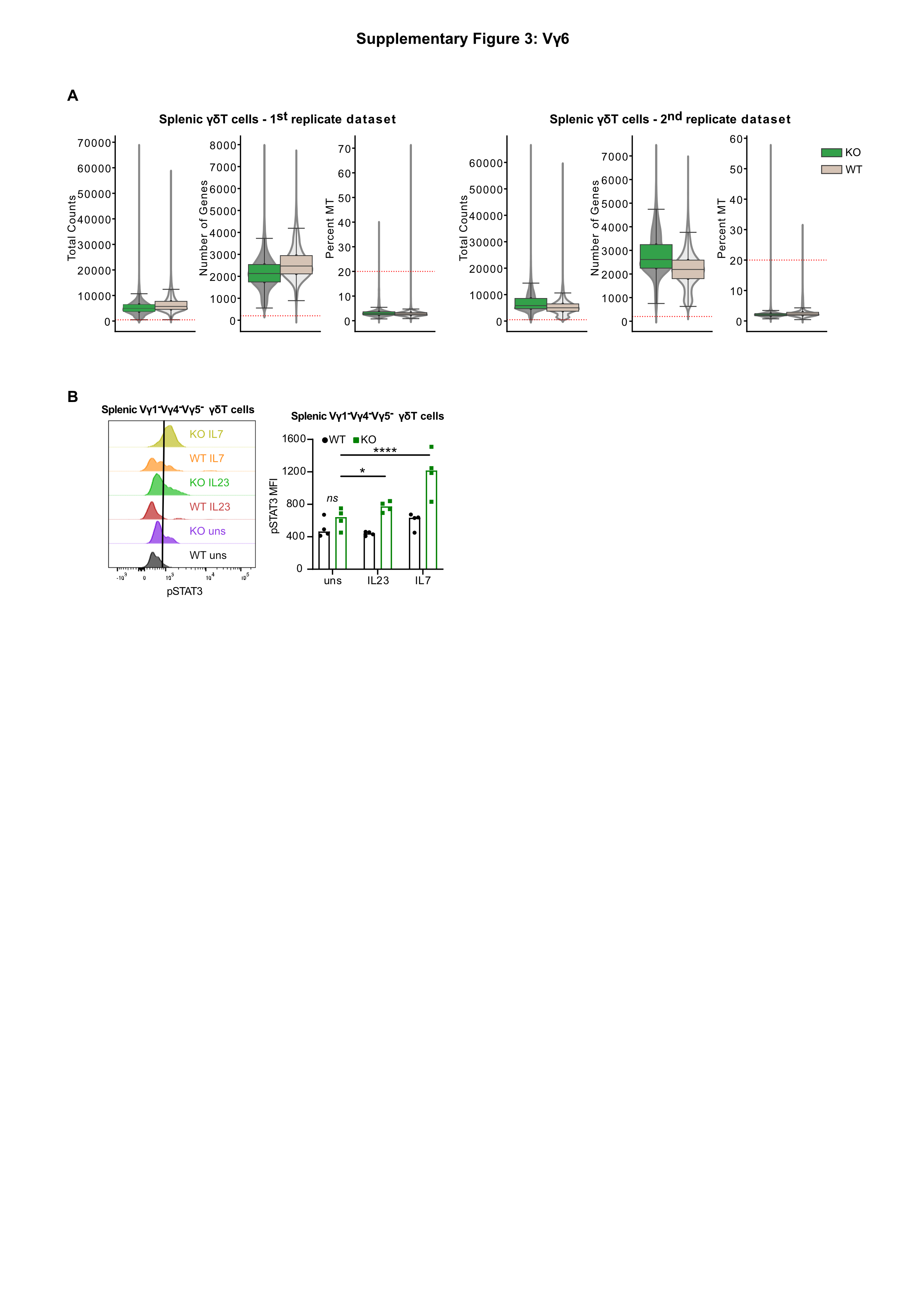

### Suppl. Fig. 4

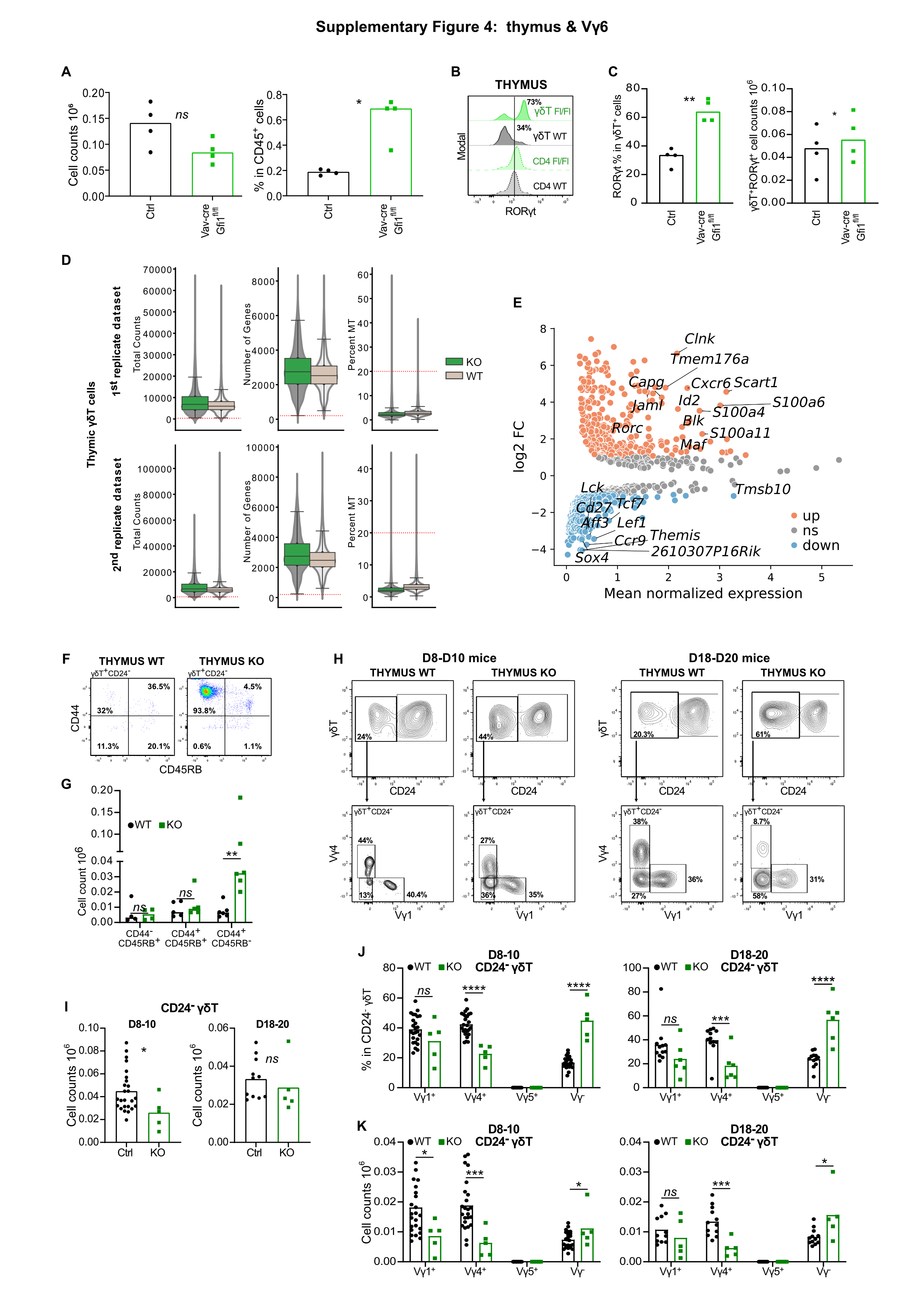

### Suppl. Fig. 5

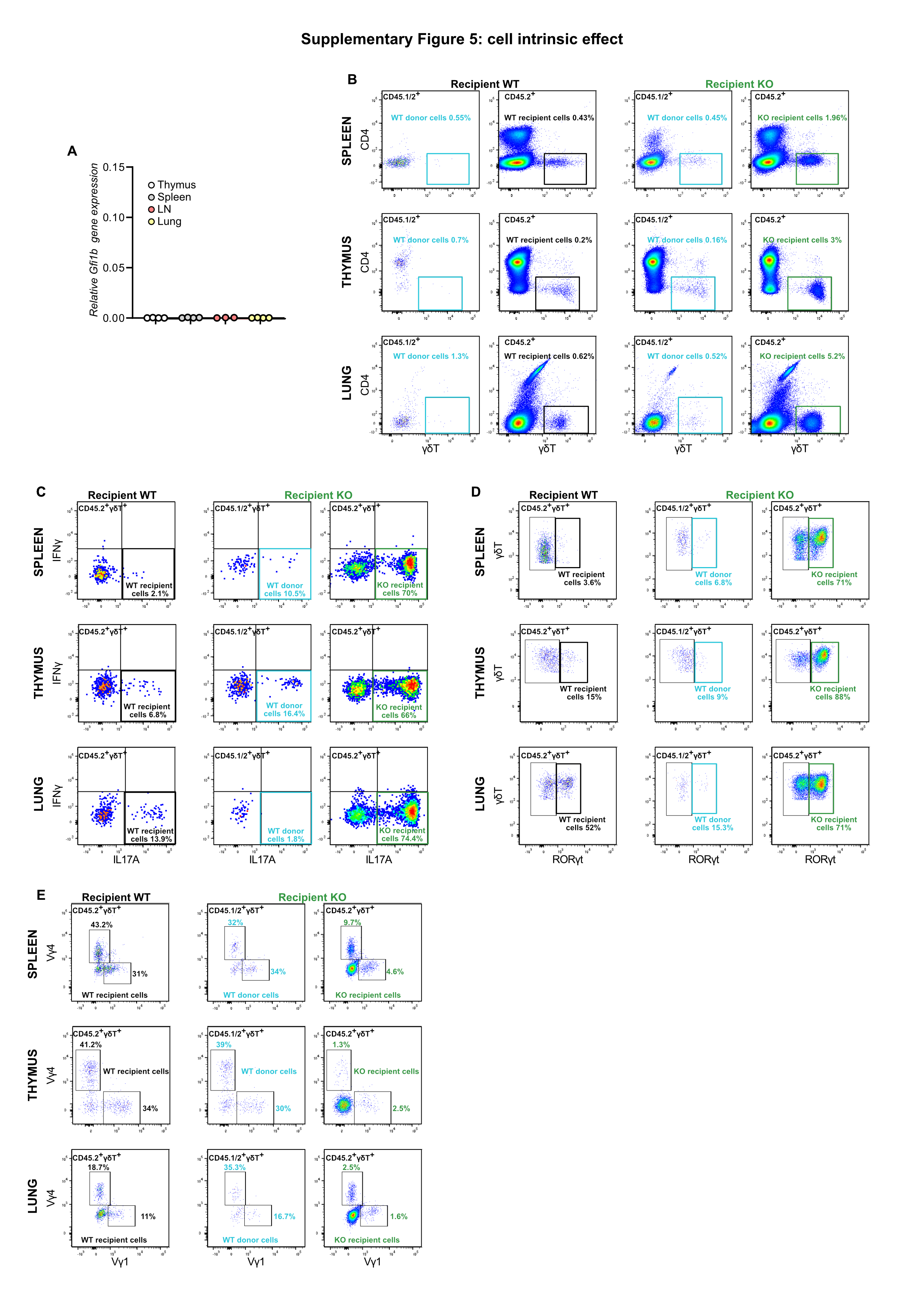

### Suppl. Fig. 6

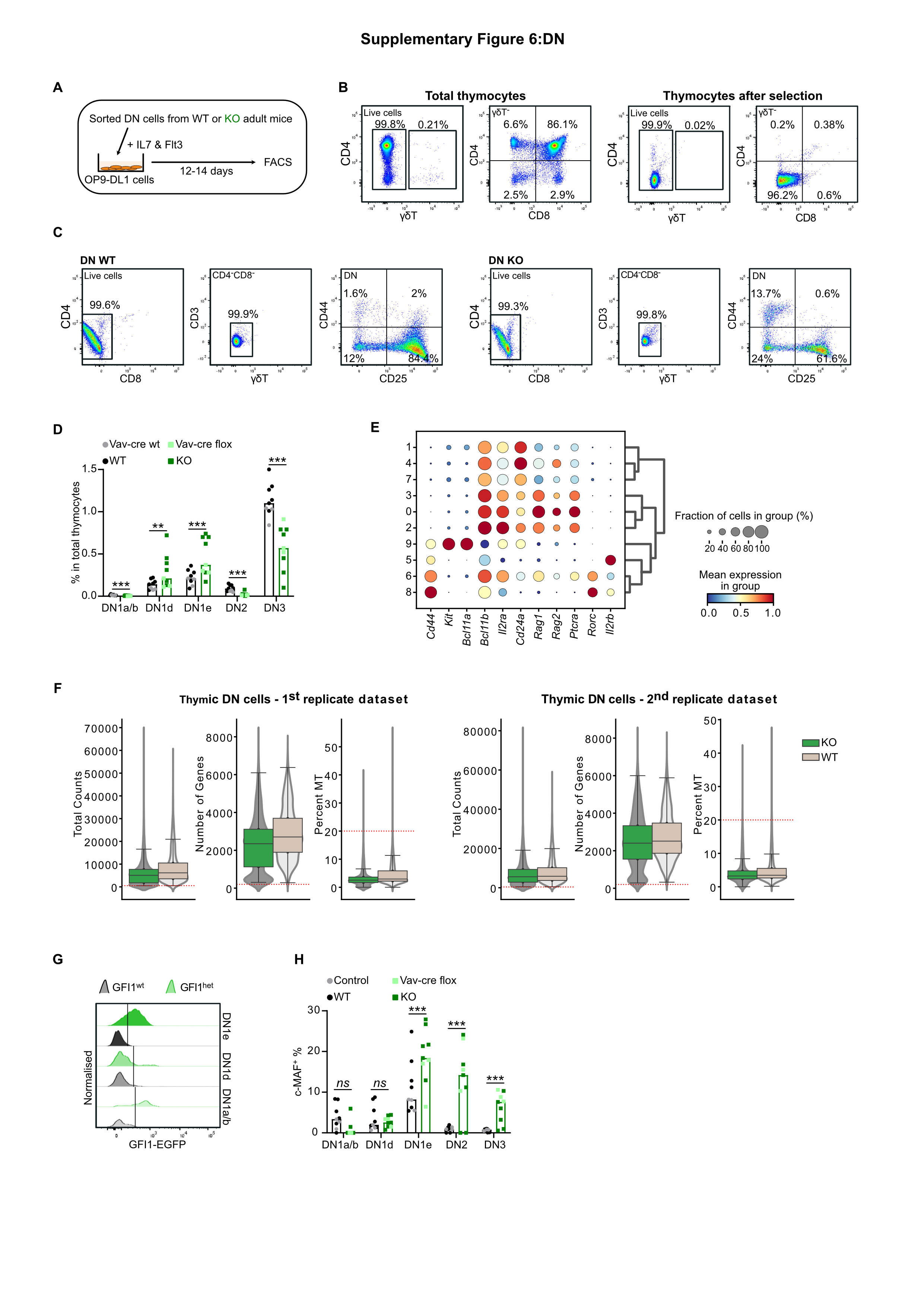

### Suppl. Fig. 7

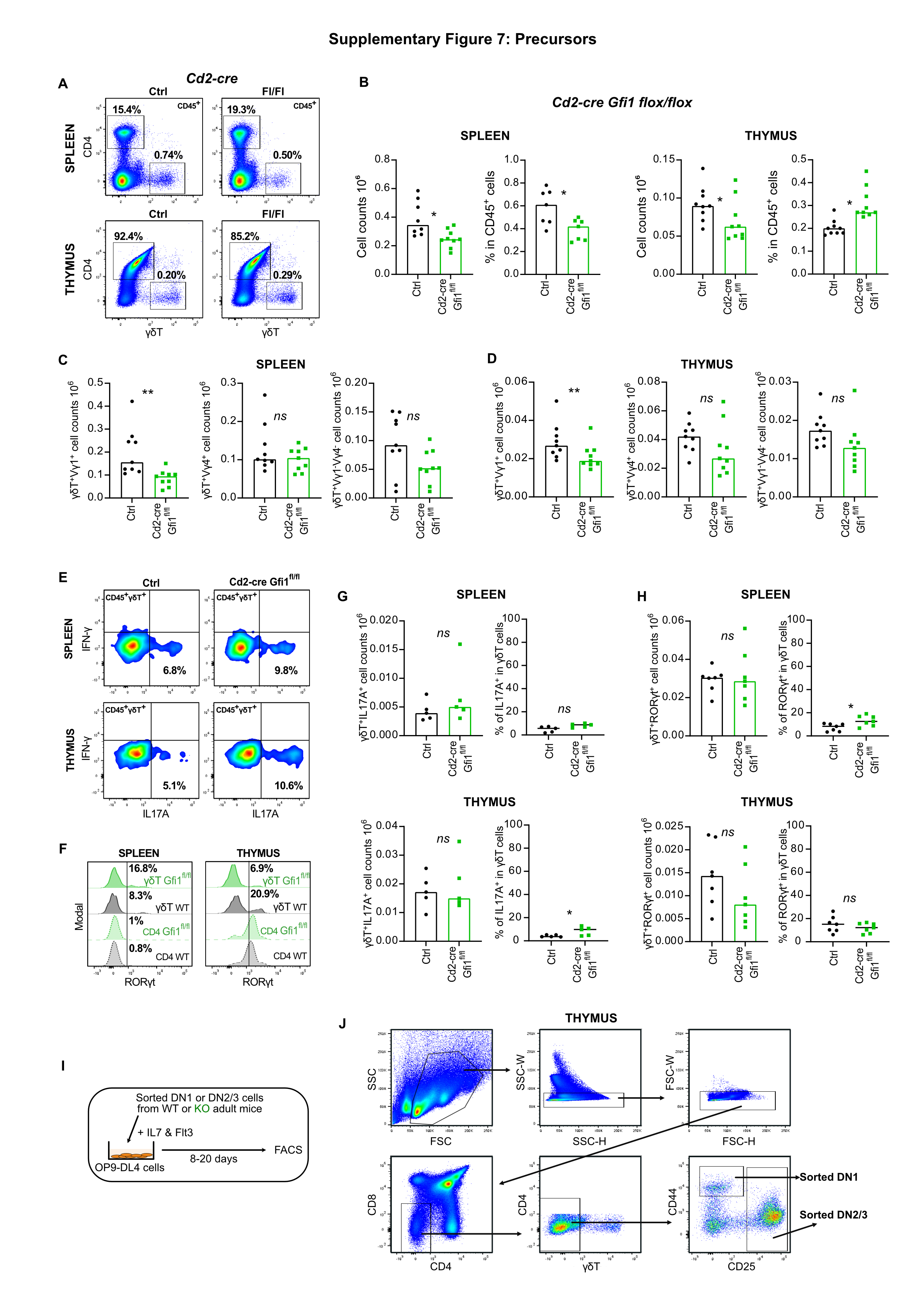

### Suppl. Fig. 8

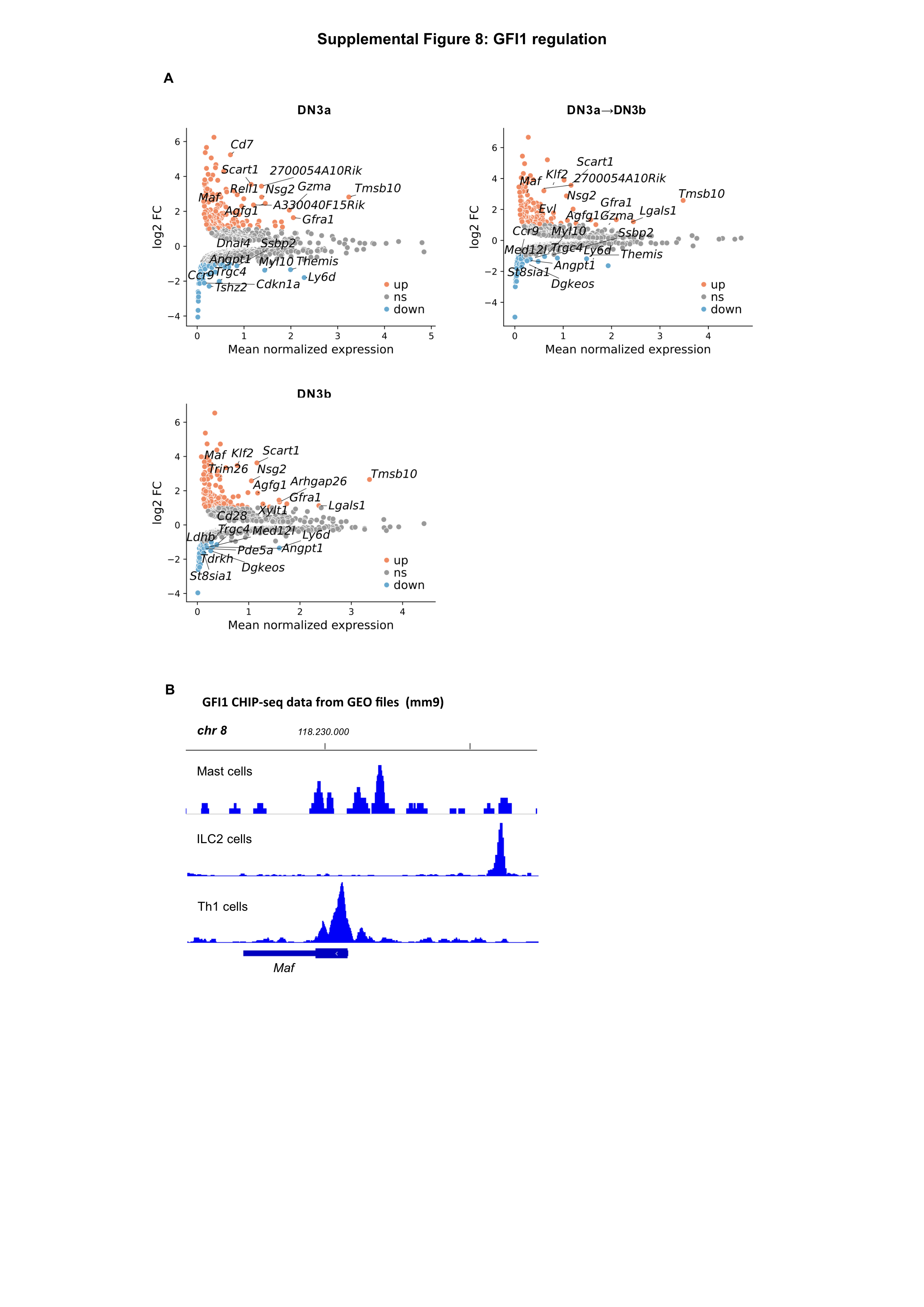
